# Objects, Faces, and Spaces: Organizational Principles of Visual Object Perception as Evidenced by Individual Differences in Behavior

**DOI:** 10.1101/2023.10.01.560383

**Authors:** Heida Maria Sigurdardottir, Inga María Ólafsdóttir

**Affiliations:** Department of Psychology, University of Iceland, Reykjavik, Iceland; Department of Psychology, Reykjavik University, Reykjavik, Iceland

**Keywords:** Object perception, face perception, high-level vision, foraging, neural networks, domain specificity, domain generality

## Abstract

What are the diagnostic dimensions on which objects differ visually? We constructed a two-dimensional object space based on such attributes captured by a deep convolutional neural network. These attributes can be approximated as stubby/spiky and animate-/inanimate-looking. If object space contributes to human visual cognition, this should have a measurable effect on object discrimination abilities. We administered an object foraging task to a large, diverse sample (N=511). We focused on the stubby animate-looking “face quadrant” of object space given known variations in face discrimination abilities. Stimuli were picked out of tens of thousands of images to either match or not match with the coordinates of faces in object space. Results show that individual differences in face perception can to a large part be explained by variability in general object perception abilities (o-factor). However, additional variability in face processing can be attributed to visual similarity with faces as captured by dimensions of object space; people who struggle with telling apart faces also have difficulties with discriminating other objects with the same object space attributes. This study is consistent with a contribution of object space to human visual cognition.

**Public Significance Statement:** The study emphasizes individual differences in visual cognition, a relatively neglected field of research. Unlike differences in other cognitive traits (e.g., Big Five personality traits, g-factor of general intelligence), we have limited knowledge on how people differ in their object processing capacity, and whether such abilities are fractionated or unitary. In this study, we ask whether visual object perception abilities are organized around an object space as evidenced by individual differences in behavior.

---

For decades, people have debated how the brain represents visual objects (Logothetis & Sheinberg, 1996; Peissig & Tarr, 2007). In anterior regions of the ventral visual pathway, which supports objects discrimination and recognition (Logothetis & Sheinberg, 1996; Milner & Goodale, 2006; Ungerleider et al., 1982; Ungerleider & Haxby, 1994), neurons can show apparently idiosyncratic activation patterns, responding vigorously to only some objects (Woloszyn & Sheinberg, 2012), but what they have in common may not be apparent (but see e.g. Tanaka, 1996). Some large-scale organization of the primate ventral stream is still known, where patches of cortex respond greatly to particular categories such as faces (Bao et al., 2020; Grill-Spector & Weiner, 2014; Kanwisher et al., 1997). Consistent with this topographical organization, face recognition deficits and exceptional abilities have been documented, but its specificity is highly debated (Barton, 2020; Barton et al., 2019; Behrmann & Plaut, 2013; Bukach et al., 2006; Burns et al., 2019; Burns & Bukach, 2021, 2022; Campbell & Tanaka, 2018; de Gelder & Van den Stock, 2018; Eimer, 2018; Farah, 2004; Farah et al., 1995; Garrido et al., 2018; Gauthier et al., 1999; Gerlach et al., 2018; Gerlach & Starrfelt, 2021, 2022; Geskin & Behrmann, 2020; Gray & Cook, 2018; Hills et al., 2015; Kanwisher, 2000; Kanwisher et al., 1997; Nestor, 2018; Plaut & Behrmann, 2013; Ramon, 2018; Robotham & Starrfelt, 2017; Rosenthal & Avidan, 2018; Rossion, 2018; Rubino et al., 2016; Starrfelt & Robotham, 2018; Susilo, 2018; Susilo & Duchaine, 2013; Susilo et al., 2015; Tarr & Gauthier, 2000; Towler & Tree, 2018).

Supposedly category-selective regions also respond to other categories (Behrmann & Plaut, 2013; Burns, Arnold, & Bukach, 2019; Gauthier, Skudlarski, Gore, & Anderson, 2000; Sigurdardottir & Gauthier, 2015; Starrfelt & Gerlach, 2007). Even individual face cells (Gross, Rocha-Miranda, & Bender, 1972) – neurons that respond particularly well to face images – respond to other objects, and both their face and object responses may reflect selectivity for *visual* properties (Vinken, Prince, Konkle, & Livingstone, 2023). Retainment of visual information by units that support face identity recognition is further demonstrated by computational work with deep convolutional neural networks trained to separate faces by their identity (Parde et al., 2021). Neuronal response patterns that support face perception may not correspond to specific semantic features.

Individual differences in object perception may also not be supported by independent abilities for different categories. A domain-general object recognition ability may account for subordinate-level object discrimination performance across categories with widely different appearance, familiarity, and meaning (e.g. Richler et al., 2019; Sunday et al., 2022; for a recent review see (Gauthier, Cha, & Chang, 2022). We will refer to this ability as *o* (Richler et al., 2019), but note that a general factor in the visual domain has also been referred to as *VG* (Hendel, Starrfelt, & Gerlach, 2019). A visual *o*-factor may explain a large part of individual differences in object perception, is at least partially modality-specific (Chow, Palmeri, & Gauthier, 2024) but see (Chow, Palmeri, Pluck, & Gauthier, 2023), and appears to be separable from low-level visual abilities as well as other cognitive abilities such as general intelligence (Chow et al., 2023; Richler et al., 2019; Richler, Wilmer, & Gauthier, 2017). Individual differences in *o* are related to neural measures of shape selectivity, including in suggested category-selective regions of the ventral visual pathway (McGugin, Sunday, & Gauthier, 2023). In the end, the visual brain might not “care” about our easily labelled or verbalized object categories.

However, it remains a fact that different parts of the ventral visual pathway do respond somewhat differently to objects of different types. While this could be based on the objects’ category, a recent paper provides strong evidence for the idea that the high-level ventral stream in non-human primates is anatomically organized not around categories, but according to the first two axes of *object space* – which dimensions can be extracted from a deep neural network trained on image classification (Bao et al., 2020; but see also Yao et al., 2023). From here on, we will use the term object space as referring specifically to this two-dimensional space. The dimensions are hard to label but may be approximated by how animate or inanimate objects appear to be, and how stubby or spiky they are. Categories such as faces may evoke activity in specific regions of the ventral visual pathway due to their *visual* qualities, as almost all human faces fall within the stubby animate-looking quadrant of object space, also known as the face quadrant. Other objects that share these qualities also evoke activity in the same brain regions of non-human primates (Bao et al., 2020).

Evidence for a visual object space in the human visual system is scarce. Functional neuroimaging studies that specifically looked for its existence produced contradictory results (Coggan & Tong, 2023; Yao et al., 2023; Yargholi & Op de Beeck, 2023). In the current preregistered study (Sigurdardottir & Ólafsdóttir, 2022, May 4; https://osf.io/q5ne8), we take a completely different approach. By measuring individual differences in object perception, we can estimate to which degree they reflect common general abilities and whether they additionally reflect an organization according to an object space.

We reasoned that if object space contributes to human visual cognition, and the quadrants of object space are at least partially anatomically and functionally separable, this should have a measurable effect on peoplés object discrimination abilities. We focused on the stubby animate-looking face quadrant of object space for two reasons. First, a lively debate has revolved around whether abilities can be specific to faces and faces arguably have the greatest claim to domain-specificity compared to any other object category; we therefore consider faces as a particularly strong test case for our hypotheses. Second, and more importantly for our purposes, there are considerable individual differences in the ability to discriminate faces (e.g., Geskin & Behrmann, 2020; Ramon et al., 2019; Russell et al., 2009; Wilmer, 2017). As people have at least some insights into their face perception abilities (Livingston & Shah, 2018; Matsuyoshi & Watanabe, 2021; Palermo et al., 2017; Shah, Gaule, Sowden, Bird, & Cook, 2015; Ventura, Livingston, & Shah, 2018), this allowed us to especially gather interest in our study of people who claim to be on the opposite sides of the spectrum of face processing ability. Sampling a wide range of abilities in our primary dependent variable (here: face discrimination ability) is critical to our experimental design. We hypothesized that face discrimination ability would be particularly associated with the ability to discriminate novel stubby animate-looking objects (preregistered hypothesis 1) and familiar stubby animate-looking objects (preregistered hypothesis 2). We separate novel and familiar objects as their association with faces may differ; while novel objects primarily have visual qualities, real objects differ in several ways, including appearance, semantics, and people’s experience with them, and these are often hard to tease apart.

## Method

### Transparency and Openness

The study followed a protocol (SHV2021-001) reviewed by the Science Ethics Committee of Icelandic Universities. Study design, data collection, and data analysis were preregistered on the Open Science Framework (Sigurdardottir & Ólafsdóttir, 2022, May 4; https://osf.io/q5ne8). All discrepancies from the plan in the preregistration are clearly stated in the manuscript. Data and analysis code are publicly available (Sigurdardottir & Ólafsdóttir, 2022, January 2; https://osf.io/2jn6c).

This paper is on object space. We also preregistered hypotheses on so-called face space (Sigurdardottir & Ólafsdóttir, 2022, May 4; https://osf.io/q5ne8), but these depended on there being measurable stable individual differences in the usage of face space. We found no such reliable individual differences, making our hypotheses regarding face space untestable. Please note that this may be restricted to face space as defined in our preregistration, as there could be stable individual differences in the usage of other types of multidimensional face spaces (see e.g., Furl et al., 2002; O’Toole, Castillo, Parde, Hill, & Chellappa, 2018; Valentine 1991; Valentine & Endo, 1992).

We report how we determined our sample size, all data exclusions, all manipulations, and all measures in the study, and the study follows JARS (Appelbaum, et al., 2018). All data, analysis code, and research materials are available (https://osf.io/2jn6c). Data were analyzed using R (R Core Team, 2020).

### Method Overview

We start with a general high-level overview of our method that we hope is accessible to all readers. Further details are found in other subsections below. We took images of various things (large and small animals and human-made objects; Konkle & Caramazza, 2013; see Method: Images) and ran them through a convolutional neural network pretrained on image classification (Mehrer et al., 2017; Mehrer et al., 2021; Muttenthaler & Hebart, 2021; Simonyan & Zisserman, 2014) and extracted layer activations of a relatively deep layer (see Method: Neural Network Activation). We chose this deep layer as it may extract complex stimulus features similar to those represented in more anterior regions of the ventral visual pathway (Bao et al., 2020; Güçlü & van Gerven, 2015). We then performed principal components analysis on these layer activations and extracted the first two principal components that explained the greatest amount of variance in the appearance of these reference objects. This allowed us to construct a two-dimensional object space that captures the two main dimensions on which the objects visually differ from one another (figure 1; see Method: Defining an Object Space). We then took tens of thousands of other object images, not used to construct object space, and projected them onto this space (see Method: Finding and Matching Stimuli). Knowing the location of each image in object space, we could pick objects, both novel and familiar, from all quadrants of object space (stubby animate-looking, spiky animate-looking, stubby inanimate-looking, and spiky inanimate-looking). Our stimulus set included novel objects from each quadrant, previously used to anatomically map object space in non-human primates (Bao et al., 2020). For each novel stubby animate-looking object, we found images of faces with closely matched object space coordinates. We also found matching stubby animate-looking familiar objects (birds, hands, seashells, bowties). We did the same for the other three quadrants. Therefore, our stimulus set also included spiky animate-looking familiar objects (birds, hands), stubby inanimate-looking familiar objects (mattresses, buckets), and spiky inanimate-looking familiar objects (hangers, exercise equipment; figures 2 and 3).

**Figure 1.**
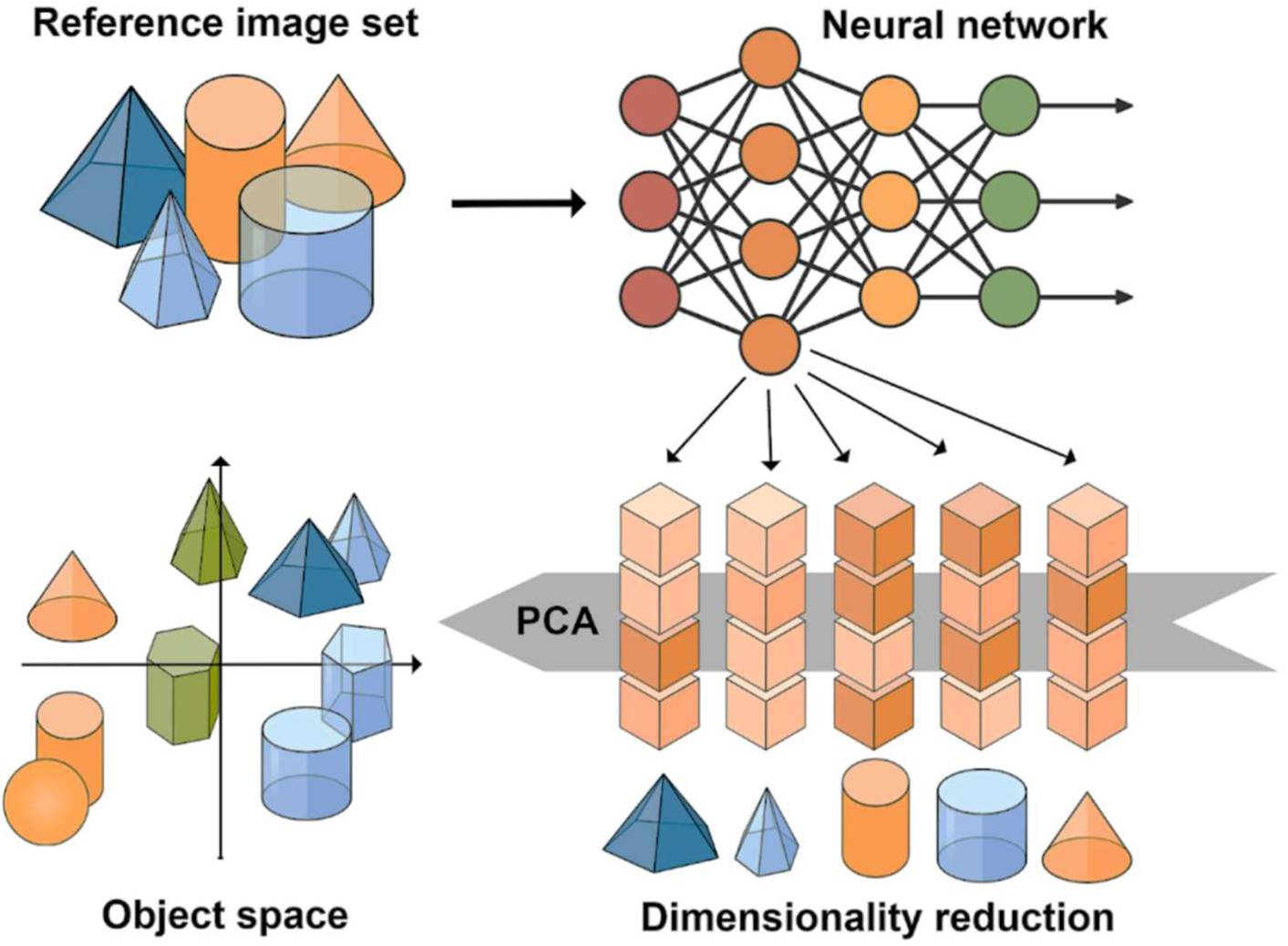
Defining object space. *Note:* The image demonstrates our overall approach to defining a visual object space. The depicted objects, network structure, activation patterns, and object space are purely illustrative. A convolutional neural network (VGG-16bn; Simonyan & Zisserman, 2014, see also Muttenthaler & Hebart, 2021) pretrained on ecologically relevant image categories (ecoset; Mehrer et al., 2017; Mehrer et al., 2021) was used to extract deep layer activations for a reference image set (large and small animate and inanimate objects; Konkle & Caramazza, 2013). Principal components analysis (PCA) on these activations was used to define an object space that should capture the main two dimensions on which the reference objects vary in appearance. Network activation patterns of other objects, not used to construct object space, were then projected to this object space to extract their coordinates. A subset of these objects was used in the experiment.

**Figure 2.**
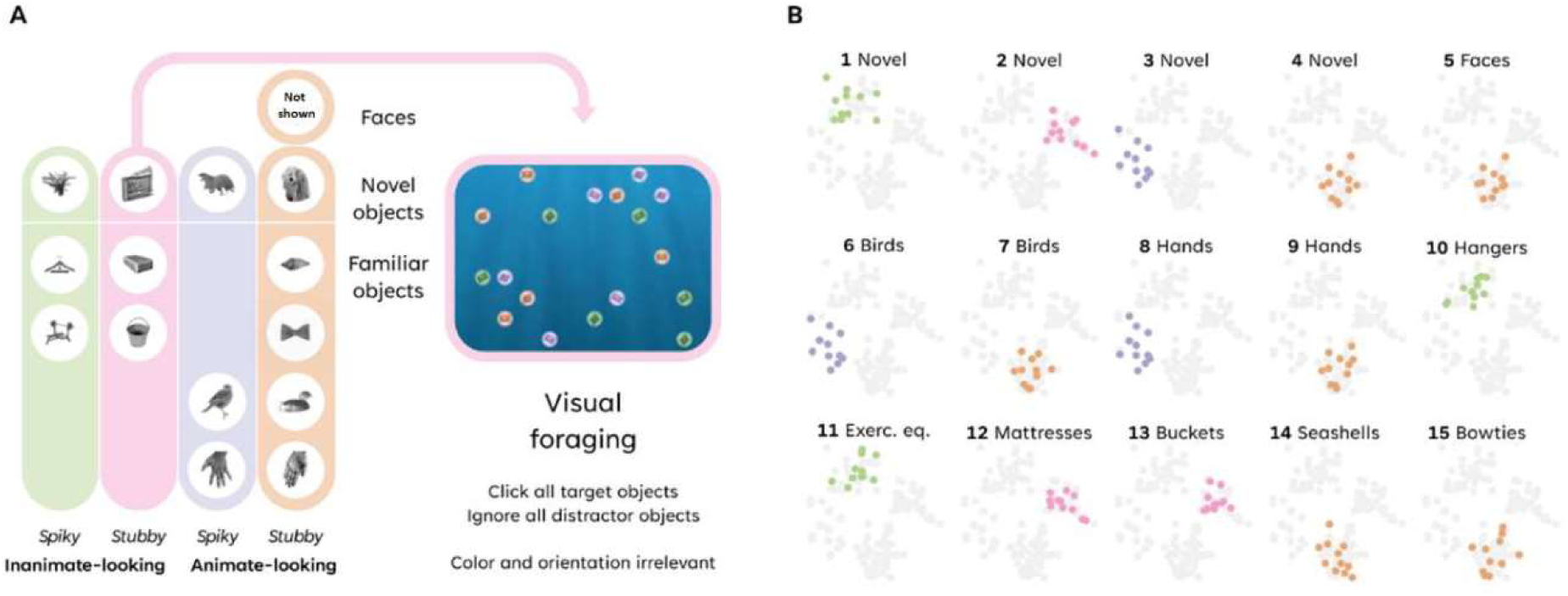
Visual foraging stimuli and their object space coordinates. *Note.* **A.** Stimuli belonged to one of four quadrants of object space. They were used in a visual foraging task where people searched for targets among distractors. **B.** Object space coordinates of stimuli (11 exemplars of each type). Quadrants of object space are represented by different colors (green: spiky inanimate-looking; pink: stubby inanimate-looking; purple: spiky animate-looking; orange: stubby animate-looking; gray: all objects). Each dot represents object space coordinates of a particular exemplar object image. An example face is not shown due to bioRxiv policy of avoiding the inclusion of photographs.

**Figure 3.**
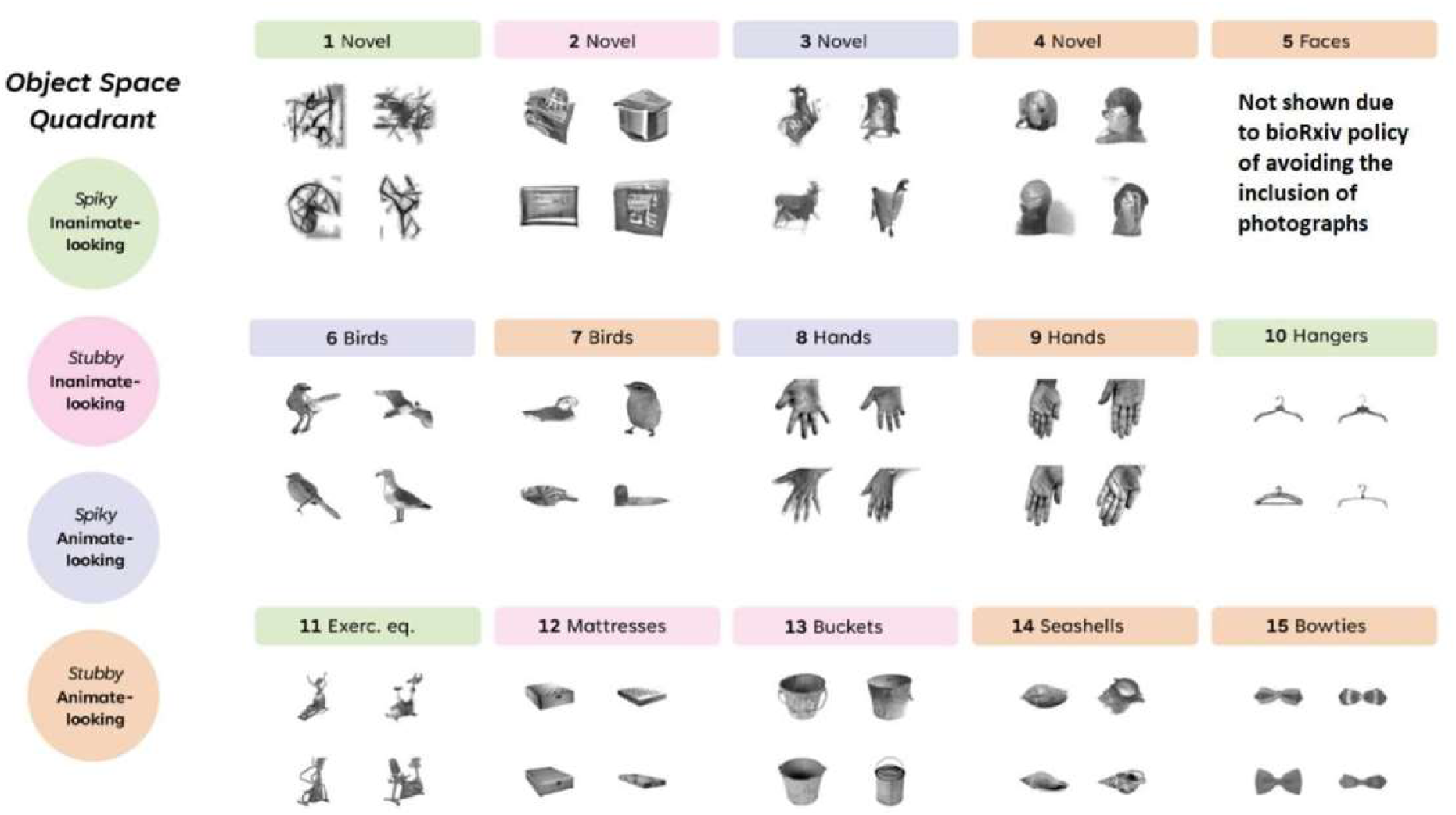
Example objects. *Note:* Example stimuli of 15 object types are shown. Quadrants of object space are represented by different colors (orange: stubby animate-looking; purple: spiky animate-looking; pink: stubby inanimate-looking; green: spiky inanimate-looking; gray: all objects).

These stimuli were then used in a visual foraging task (figure 2) where a large sample of participants (N = 511) with greatly varying face processing abilities had to search for targets among distractors. On each experimental trial, a sample stimulus was briefly shown at screen center, followed by the appearance of multiple targets identical to the sample, and multiple distractors that depicted another exemplar of the same object type. Participants were asked to click on all targets as fast as possible and avoid distractors. Color overlay and orientation of targets and distractors varied to discourage low-level visual matching strategies. The experiment also included control trials without objects where people searched for black targets among white distractors, or white targets among black distractors. Discrimination ability was then defined for each trial type. A person’s face discrimination ability was therefore defined as their foraging speed on face trials, bucket discrimination ability was foraging speed on bucket trials, and so on. We then assessed whether individual differences in face discrimination depended on their location in visual object space.

### Participants

Participants were unpaid volunteers recruited by word of mouth, email listservs, social media, and other media. We especially encouraged those with self-perceived face recognition abilities on opposite sides of the spectrum to participate, ranging from poor to excellent face recognizers, but participation was open to people regardless of abilities in face recognition. The study was run online, and participation was anonymous. Participants could stop at any time by simply closing their browser without any consequences to them, and we did not use their data in the current study unless they finished all tasks. A total of 514 people completed all tasks. Thereof, three participants had mean targets per second at least three standard deviations away from the sample mean in visual foraging. These data were excluded from further analysis, leaving us with a final sample of 511 participants. Participants were asked about their gender, age, and educational level, choosing between the following options: 357 were women, 144 were men, and 10 were non-binary, other gender, or chose not to answer. Twenty-five people were between the ages of 18-24, 86 were 25-34 years, 134 were 35-44 years, 132 were 45-54 years, and 134 were 55 years of age or older. A total of 26 participants had completed primary education or less, 122 had completed secondary education, 156 undergraduate education, and 207 graduate education. All participants reported normal or corrected-to-normal vision and gave informed consent. For sample size rationale, stopping rule for data collection, and inclusion/exclusion criteria, see preregistration (Sigurdardottir & Ólafsdóttir, 2022, May 4; https://osf.io/q5ne8).

### Procedure

The study was run online through Pavlovia (https://pavlovia.org/) using PsychoPy (Peirce et al., 2019; Peirce, 2007). Participants first completed a short speed click task where they clicked on several consecutive targets. Those who did this slowly were reminded to use a computer mouse instead of a touchpad and had to repeat the task before being allowed to continue. Participants then answered background questions on their age bracket, gender, and educational level. Next, people completed the 20-item prosopagnosia index (PI20; (Shah et al., 2015), Icelandic version), a self-report on problems with faces, and the Cambridge Face Memory Test (CFMT; (Duchaine & Nakayama, 2006) which measures people’s memory for unfamiliar faces. Finally, participants completed a visual foraging task where they searched for target objects among distractor objects. Participants were given feedback on their general performance upon study completion.

### Visual Foraging

In this main experimental task, people searched for faces as well as other familiar and novel objects. These objects either shared or did not share visual qualities with faces as objectively defined by their coordinates in a visual object space derived from activation patterns in a deep layer of a neural network trained on object classification. We will refer to the dimensions of object space as animate-inanimate-looking and stubby-spiky-looking, following Bao et al. (Bao et al., 2020), but note that these are only approximate labels for these main dimensions on which objects differ visually.

The foraging task included 11 trials from each of the following 17 conditions (187 trials per person): seven stubby animate-looking object types (novel objects, hands, birds, bowties, seashells, faces matched to novel objects, faces matched to each other), three spiky animate-looking object types (novel objects, hands, birds), three spiky inanimate-looking object types (novel objects, exercise equipment, hangers), three stubby inanimate-looking object types (novel objects, buckets, mattresses), and a feature foraging control condition. Each object image appeared on two trials in the experiment, once as a target and once as a distractor. Each target-distractor pair only appeared once, so if image A was a target and B a distractor in one trial, then B was a target and not-A a distractor in another trial. Each trial in the visual foraging task consisted of a sample stimulus shown at screen center for 1 second followed by 9 targets that were identical to the sample and 9 distractors that showed another object not identical to the sample (figure 2). Targets and distractors within a trial always belonged to the same condition. In all but control trials, stimulus orientation and color of foraging items was varied to discourage the use of low-level visual strategies. Participants foraged for targets as fast as they could by clicking each of them with the mouse. Targets disappeared as they were clicked; distractors jiggled and stayed on-screen. The trial ended when all targets had been clicked or when 10 seconds had passed from the onset of the foraging items, whichever came first. A trial was followed by feedback where 0 to 9 stars were shown depending on the number of targets clicked.

### Images

#### Image manipulation

All objects were stripped of their background (if they had one) so that only a single object was shown on a blank background. To standardize size and borders, images were processed in IrfanView (https://www.irfanview.com). All images were cropped and then resized so that the longer side of each object (vertical or horizontal) was the same size as the longer side of all other objects (bounding box longer side 680 pixels). We then added a square, white background to all images, before we resized them again to 224 x 224 pixels. All images were therefore square, equally sized, with an equally wide, white border around them. Images were changed to grayscale and run through the SHINE Toolbox (Willenbockel et al., 2010) to equate mean brightness and contrast.

#### Reference set

We defined object space using a reference set of 120 animals and 120 inanimate objects with large and small real-world size, which were selected to have broad coverage over the categories (image courtesy Konkle Lab: http://konklab.fas.harvard.edu/; Konkle & Caramazza, 2013). Subsets of these objects map onto different anatomical zones of the human ventral stream (Konkle & Caramazza, 2013).

#### Stimuli

Images for the foraging study were chosen from five additional image sets that contained over 30,000 images of various objects. A set of 80 novel objects (images provided by the Tsao Lab: https://tsaolab.berkeley.edu/), were projected onto the object space. The images originally formed four groups of 20 items that mapped onto the four quadrants of object space (stubby-animate, stubby-inanimate, spiky-animate, and spiky-inanimate) in Bao et al. (2020) their extended data fig. 6) and evoked preferential activity in different networks of the macaque inferotemporal cortex. A set of 4760 images of various familiar objects (image credit: the University of California San Diego (UCSD) Vision and Memory Lab: https://bradylab.ucsd.edu/stimuli.html; Konkle et al., 2010) were also projected onto our object space. An image set of 3362 Caucasian faces of 106 individuals (image credit: Michael J. Tarr, Carnegie Mellon University, http://www.tarrlab.org/ – funding provided by NSF award 0339122 (Righi et al., 2012)) was projected onto our object space. Additionally, we used images from 11k Hands (image courtesy Mahmoud Afifi: https://sites.google.com/view/11khands; Afifi, 2019), and Caltech-UCSD Birds-200-2011 (image courtesy Caltech Vision / UCSD Vision: http://www.vision.caltech.edu/visipedia/CUB-200-2011.html; Wah et al., 2011).

### Neural Network Activation

All images were run through the convolutional neural network VGG-16bn (Simonyan & Zisserman, 2014) trained on the ecoset image set (Mehrer et al., 2021) using batch normalization (code implemented through the Python toolbox THINGSvision (Muttenthaler & Hebart, 2021) https://github.com/ViCCo-Group/THINGSvision). The activation from the second fully connected (fc) layer corresponding to each image was extracted – a vector of length 4096, with each vector item corresponding to the activation of a particular node.

### Defining an Object Space

The layer activations corresponding to the reference set were run though Principal Component Analysis (PCA). The first two principal components were extracted; they explained a total of 20.1% of variance in the original 4096 variables.

### Finding and Matching Stimuli

The four types of novel objects (that looked stubby-animate, stubby-inanimate, spiky-animate, and spiky-inanimate; Bao et al., 2020) were projected onto our object space. Eleven images from each type were selected to minimize overlap in object space location between images from different types. Images from four other image sets were then projected onto object space and for each novel object image, an image was found that projected onto the same or very similar location in object space. These images were therefore matched in visual qualities on dimensions of object space. Objects from six categories of familiar inanimate objects were matched with the novel objects: bowties and seashells with stubby animate-looking novel objects, mattresses and buckets with stubby inanimate-looking novel objects, and hangers and exercise equipment with spiky inanimate-looking novel objects. We chose two familiar inanimate categories per quadrant by automatically (i.e., via scripting) picking the 11 objects from each category from the familiar objects image set (image credit: the University of California San Diego (UCSD) Vision and Memory Lab: https://bradylab.ucsd.edu/stimuli.html; Konkle et al., 2010) that best matched the coordinates of the novel objects of each quadrant, and then manually going over each category by plotting the object space coordinates of the best 11 objects and picking familiar inanimate categories that coordinate-matched well and did not bleed into other quadrants. Few familiar inanimate objects projected to the spiky animate-looking quadrant of object space, and we therefore could not match them to corresponding novel objects. Hands and birds clustered in the animate-looking part of object space and were matched with animate-looking stubby and spiky novel objects via automatic scripting; no good matches for hands or birds were found for inanimate-looking novel objects. Exclusion criteria for hand images were jewelry or salient sleeves. If two images that matched with novel objects were clearly of the same hand, the image with an object space distance further from its assigned novel object was excluded and a new match found. Exclusion criteria for bird images were more than one bird in each image. As expected, most faces projected to the stubby-animate quadrant (face quadrant) of object space and were therefore only matched with stubby animate-looking novel objects (object-matched faces). Exclusion criteria for faces were as follows: sunglasses or obvious wigs. If two images of the same person were matched with different novel objects, the image with a distance in object space further from its assigned novel object was excluded and a new match found. In addition to these 15 types of target-distractor pairs, we included difficult face foraging trials where we handpicked 11 images of men that had similar facial features, hairstyles, and expressions, and were all looking slightly to the left (self-matched faces). Lastly, we included 11 feature foraging control trials where participants searched for black discs among white discs and vice versa. 17 types of target-distractor pairs were therefore shown in the experiment: four types of novel objects, six types of familiar inanimate objects, two types of birds, two types of hands, two types of faces, and feature foraging control stimuli.

### Target-Distractor Pairs

The novel objects were randomly paired together within each object space quadrant (stubby-animate, stubby-inanimate, spiky-animate, spiky-inanimate) as targets and distractors. Mean target-distractor distance was not significantly different between the four quadrants. Control trials consisted of black targets and white distractors or white targets and black distractors. For each of the other conditions, the target-distractor pairs were matched with the novel object pairs. For instance, if hand A was matched with novel object A and hand B with novel object B, then hand A and hand B were selected as a target-distractor pair, because novel objects A and B were paired together. Across the experiment, there were 11 unique target-distractor pairs per condition. The target and the distractor were separated in object space by a known distance varying from small (similar object space coordinates) to large (dissimilar object space coordinates). This distance should vary in a similar way for each experimental condition. Faces from the face quadrant of object space matched to each other (self-matched faces) are an exception; they were not chosen to match any other objects but were hand-picked by the researchers. These 11 faces formed a tight cluster in object space, consistent with their visual similarity. We include these faces even though they are not matched to the other objects to include particularly challenging face foraging trials where cues unrelated to facial identity (e.g., viewpoint) are minimized.

### Data Analysis

Target-distractor visual dissimilarity in the foraging task was defined as Euclidean distance in object space. Object discrimination performance was defined as the number of targets clicked per second. For each condition, separately for each participant, we calculated the median targets per second. Descriptive statistics and reliability measures for all conditions be found in table 1 (for boxplots showing distributions, see supplementary figure s1).

**Table 1.** Descriptive statistics and reliability of foraging measures. Discrimination performance of each participant is defined as their median targets per second in a particular condition.

| *Foraging conditions* | *Mean*<br>*performance* | *SD performance* | *Cronbach's alpha* |
| --- | --- | --- | --- |
| *Control foraging* | 1.40 | 0.23 | 0.97 |
| *Novel spiky-inanimate* | 1.00 | 0.19 | 0.93 |
| *Hanger spiky-inanimate* | 1.11 | 0.17 | 0.95 |
| *Exer. eq. spiky-inanimate* | 0.87 | 0.16 | 0.90 |
| *Novel stubby-inanimate* | 1.05 | 0.18 | 0.94 |
| *Bucket stubby-inanimate* | 1.02 | 0.18 | 0.94 |
| *Mattr. stubby-inanimate* | 0.86 | 0.18 | 0.90 |
| *Novel spiky-animate* | 1.03 | 0.18 | 0.93 |
| *Bird spiky-animate* | 1.08 | 0.17 | 0.93 |
| *Hand spiky-animate* | 0.81 | 0.17 | 0.89 |
| *Novel stubby-animate* | 1.12 | 0.18 | 0.95 |
| *Bird stubby-animate* | 1.09 | 0.17 | 0.94 |
| *Hand stubby-animate* | 0.80 | 0.16 | 0.90 |
| *Bowtie stubby-animate* | 1.02 | 0.18 | 0.93 |
| *Seashell stubby-animate* | 1.08 | 0.18 | 0.94 |
| *Object-matched faces* | 1.04 | 0.17 | 0.95 |
| *Self-matched faces* | 0.81 | 0.18 | 0.93 |

Two preregistered hierarchical regressions were conducted to assess whether face foraging (object-matched faces) was associated with a) novel stubby animate-looking objects, and b) familiar stubby animate-looking objects. Demographic variables were entered in the first step and were coded as follows: Gender was coded as 1 = man, 0 = woman/non-binary/other/choose not to answer. This coding was chosen *a priori* as per our preregistration, as dummy coding as e.g. man (1 = yes; 0 = no) and woman (1 = yes; 0 = no) was deemed likely to pose a multicollinearity problem due to few participants choosing non-binary/other/choose not to answer. Age was coded as 1 = 18-24 years, 2 = 25-34 years, 3 = 35-44 years, 4 = 45-54 years, and 5 = 55 years and older. Educational level was coded as 1 = completed primary school or less, 2 = completed secondary school/gymnasium or compatible degrees, 3 = completed an undergraduate degree such as BSc, BA, BEd, and 4 = completed a graduate degree such as MA, MSc, PhD. Control foraging was added in step two of the hierarchical regressions, foraging for objects other than stubby animate-looking was added in step 3, and lastly foraging for stubby animate-looking objects was added in step 4.

For the purpose of a follow-up analysis on stubby and spiky-looking hands and birds, new indices of stubby-looking birds and hands and spiky-looking birds and hands were made. For each participant, the means of a) the median targets per second for stubby-looking hands and stubby-looking birds, and b) spiky-looking birds and spiky-looking hands were calculated.

Because the correlation between scores on the PI20 and the CFMT (−0.51) reached a preregistered threshold (−0.5), these measures were combined into a single face recognition measure. For each participant, scores from PI20 and CFMT were z-scored, the z-score of PI20 was negated (so e.g., 1.5 became −1.5 and vice versa) and the mean of these were taken to indicate a person’s face recognition ability.

For exploratory multiple regressions, new indices of mean stubby animate-looking objects and mean non-stubby animate-looking objects were calculated. Mean stubby-animate: For each participant, the mean of the median targets per second for novel stubby animate-looking objects, stubby animate-looking birds, stubby animate-looking hands, stubby animate-looking bowties, and stubby animate-looking seashells was calculated. Mean non-stubby-animate: For each participant, the mean of the median targets per second for novel spiky inanimate-looking objects, novel stubby-inanimate looking objects, novel spiky animate-looking objects, spiky animate-looking birds, spiky animate-looking hands, spiky inanimate-looking hangers, spiky inanimate-looking exercise equipment, stubby inanimate-looking buckets, and stubby inanimate-looking mattresses was calculated.

## Results

Almost all faces belong to the stubby animate-looking quadrant of object space (face quadrant). We hypothesized (preregistration: https://osf.io/q5ne8) that face foraging would be positively associated with foraging for novel stubby animate-looking objects (hypothesis 1) and with familiar stubby animate-looking objects (hypothesis 2).

### Predicting Face Discrimination

We assessed these hypotheses by performing two preregistered hierarchical regressions with face discrimination ability (object-matched faces) as the dependent variable. In both cases, demographics (age, gender, and educational level) were entered at step 1, and control foraging performance was added at step 2. At stage 1 of each model, age, gender, and education accounted for around 37% of the variance in face discrimination. Higher educational level was associated with better face discrimination ability, while advanced age was associated with poorer face discrimination ability (supplementary figures s2 and s3). Adding the control trials at stage 2 greatly improved the model which now explained around 66% of the variance in the dependent variable. Control foraging, which is moderately to highly correlated with all other foraging measures (range: r = 0.58 to r = 0.82; for all zero-order correlations, see supplementary table s1), should capture various factors of little interest that do not directly involve object discrimination ability, and likely provides an upper limit for a person’s foraging speed.

On step 3 of the first hierarchical regression, we added three factors: foraging performance for novel spiky inanimate-looking objects, novel stubby inanimate-looking objects, and novel spiky animate-looking objects. The included variables now accounted for 82% of the variance in face discrimination ability. Finally, and importantly for hypothesis 1, adding foraging performance for novel stubby animate-looking objects at stage 4 significantly (*p* = 4.95×10^−8^, two-sided) improved the model from step 3, with 83% explained variance (table 2).

**Table 2.**
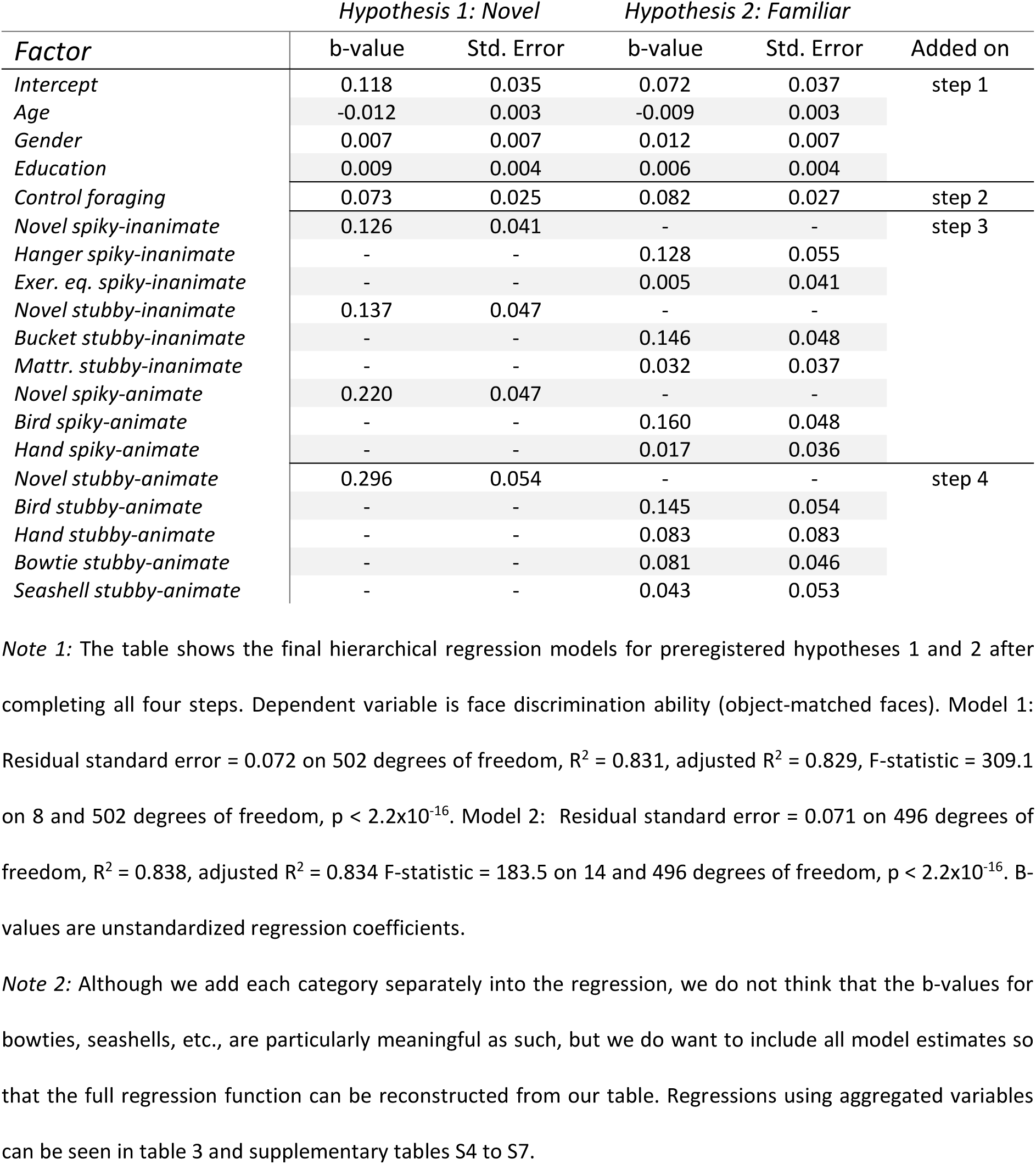
Association between face discrimination (object-matched faces) and discrimination of stubby animate-looking objects.

On step 3 of the second hierarchical regression, we added six factors: foraging performance for spiky animate-looking birds and hands, spiky inanimate-looking hangers and exercise equipment, and stubby inanimate-looking buckets and mattresses. In step 4 of this second regression, we added four factors: foraging performance for stubby animate-looking birds, hands, bowties, and seashells. This again greatly improved the model from that of stage 2, where the included factors explained a total of 83% of the variance in face foraging. Importantly for hypothesis 2, adding foraging performance for familiar stubby animate-looking objects (birds, hands, bowties, seashells) further improved the model from step 3 to step 4 (p = 7.28×10^−4^, two-sided; table 2) where all included factors now explained a total of 84% of variance in face discrimination ability (table 2).

The great improvement in the models from step 2 to step 3 likely reflects individual differences in a general object perception factor *o* (Richler et al., 2019; Hendel et al., 2019) that contributes to people’s ability to discriminate all sorts of objects, including faces. Consistent with this interpretation, face foraging is highly correlated with all object foraging measures (range: r = 0.76 to r = 0.87), and all object foraging measures are themselves also highly correlated (range: r = 0.77 to r = 0.91; for all zero-order correlations, see supplementary table s1). Nonetheless, adding stubby animate-looking objects to the models at step 4 further improves their predictions for face discrimination ability. Both preregistered hypotheses 1 and 2 were confirmed (see also supplementary tables s2 and s3).

### Keeping Category Constant

The variance in face foraging (object-matched faces) uniquely explained by foraging for stubby animate-looking objects was small. This is not unexpected given that a person’s ability to forage for faces is already extremely well predicted from demographics and performance on other types of foraging trials, leaving little non-noise variance left to explain.

We nonetheless chose to take a closer look at our results. The regression coefficient for novel stubby animate-looking object foraging was larger than all other types of foraging included as independent predictors in the final model, as predicted by hypothesis 1 (table 2). The regression coefficient for novel spiky inanimate-looking objects was the smallest of all novel object foraging types, in accordance with these objects being the furthest in object space from faces. Regression coefficients for familiar object foraging showed a less clear pattern (table 2). This can be deceptive as regression coefficients reflect the *unique* contribution of a particular condition. When multiple conditions with a common factor (e.g., several different stubby animate-looking things in stage 4) are added, this common attribute of interest (e.g., “stubby-animateness”) is essentially factored out. This less clear pattern could however also be due to the difficulty of fully separating semantics and previous experience with object categories from their visual qualities.

We can however do this for two categories as we had access to over ten thousand images of birds and hands, allowing us to include both stubby animate-looking and spiky animate-looking items. We therefore followed up with exploratory analyses where we focused on unique variance in face discrimination ability (object-matched faces) explained by stubby vs. spiky familiar objects of the same category (figure 4). This showed that face discrimination was more associated with average performance for stubby animate-looking birds and hands, partialling out average performance for spiky animate-looking birds and hands (r_partial_ = 0.425), than with average performance for spiky animate-looking birds and hands, partialling out average performance for stubby animate-looking birds and hands (r_partial_ = 0.299). This is unlikely to reflect differential reliability of the measures as they both had excellent reliability (Cronbach’s alpha for stubby animate-looking birds and hands: 0.96; Cronbach’s alpha for spiky animate-looking birds and hands: 0.95). We therefore are more confident in our assessment that individual differences in face discrimination may partially tap into individual differences in the processing of stubby animate-looking objects (for further analysis, see supplementary tables s2 to s7).

**Figure 4.**
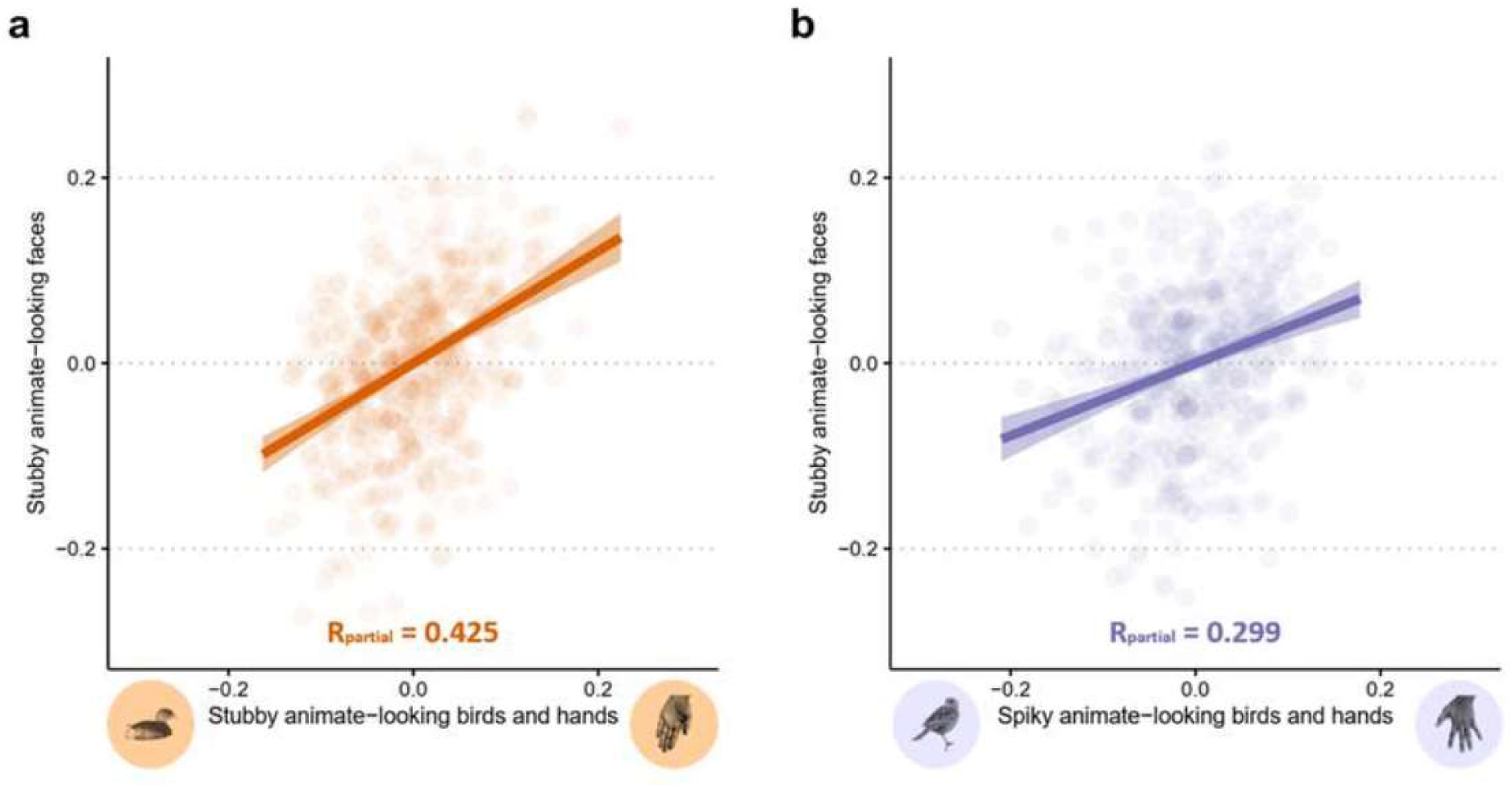
Association between face discrimination ability (object-matched faces) and stubby vs. spiky animate-looking birds and hands. *Note:* **A.** Partial correlation plot between face discrimination ability and average discrimination ability for stubby animate-looking birds and hands; axes show residuals after partialling out average discrimination ability for spiky animate-looking birds and hands. **B.** Partial correlation plot between face discrimination ability and average discrimination ability for spiky animate-looking birds and hands; axes show residuals after partialling out average discrimination ability for stubby animate-looking birds and hands (targets per second).

### Clustering of Abilities

To visualize our results for hypotheses 1 and 2, we performed multidimensional scaling on individual differences in foraging. As can be seen in figure 5, faces formed a separate cluster, indicating that people’s performance on a given face foraging trial can generally best predict their performance on another trial of the same type. Of note, the face cluster is closer to the stubby animate-looking cluster than it is to the cluster for other objects, both for novel objects (figure 5a, hypothesis 1) and familiar objects (figure 5b, hypothesis 2), indicating that foraging performance for stubby animate-looking objects more closely predicts face foraging.

**Figure 5.**
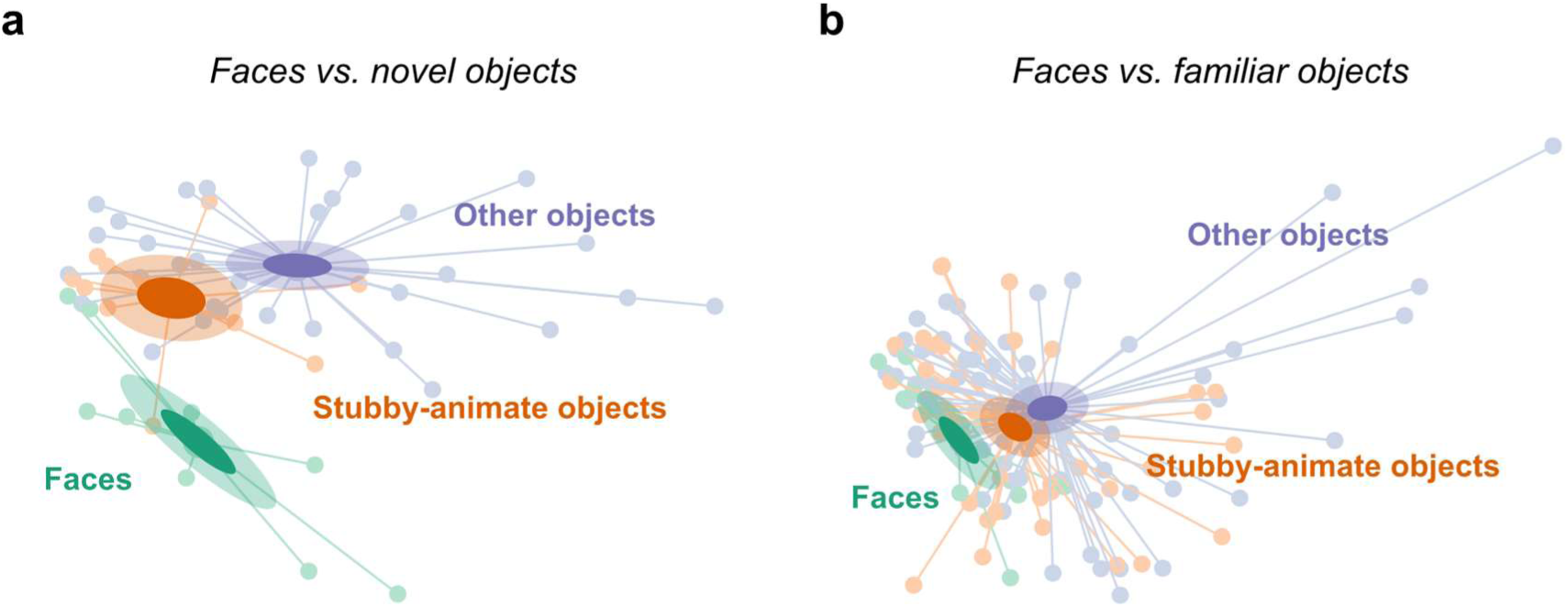
Clustering of individual differences in visual foraging. *Note:* Multidimensional scaling was performed on the correlation matrix for performance (targets per second) on individual trials. This was done separately for a) trials with novel objects and object-matched faces, and b) trials with familiar objects and object-matched faces. Each dot depicts a single trial and the distance between two points is approximately equal to their dissimilarity. Trials are clustered based on our preregistered analysis pipeline: the faces cluster corresponds to trials contributing to the face discrimination dependent variable in table 2, the stubby-animate objects cluster corresponds to trials contributing to factors added at step 4 in table 2, and the other objects cluster corresponds to trials contributing to factors added at step 3 in table 2. Dots are connected to their cluster’s barycenter via lines. Confidence ellipses for barycenters of the three clusters are shown, both for the 95% confidence level (light ellipses) and the 50% confidence level (dark ellipses).

### Predicting Face Recognition

The preregistered analysis pipeline for hypotheses 1 and 2 concerned predicting foraging performance for faces matched for object space coordinates to other stubby animate-looking objects (object-matched faces, see figure 2b). However, we assessed face processing in additional ways. We also included difficult faces foraging trials where cues unrelated to facial identity (e.g., viewpoint) were minimized. We did this by hand-picking faces of 11 male individuals looking slightly to the left with a similar expression, similar hairstyle, and no beard or glasses. These 11 faces formed a tight cluster in the stubby animate-looking quadrant of object space (face quadrant), consistent with their visual similarity. We refer to performance on these trials as self-matched face discrimination ability. Participants also filled out a questionnaire on their face recognition abilities in everyday life (20-item Prosopagnosia Index, PI20; Shah et al., 2015), and tried to recognize previously memorized faces (Cambridge Face Memory Test, CFMT; Duchaine & Nakayama, 2006). As defined in our preregistration, we combined PI20 and CFMT into a single face recognition index.

In our preregistered analysis, we predicted people’s ability to discriminate faces matched to object space coordinates of non-face objects (object-matched faces). We wanted to see if discrimination ability for stubby animate-looking objects could predict face processing more generally. We therefore took a better look at our additional measures of face processing ability. As our preregistered hypotheses held for both novel and familiar objects, we collapsed across them to make two indices (based on foraging excluding all face and control trials): discrimination ability for stubby animate-looking objects and for other objects (for additional analysis where we aggregate separately across the four quadrants of object space, see supplementary table s4). Aggregating across variables is expected to increase reliability of our measures even further and may better reveal the influence of a broader construct (Richler et al., 2019). The aggregated measures indeed had excellent reliability (Chronbach’s alpha for stubby animate-looking objects: 0.98; Chronbach’s alpha for other objects: 0.99).

In an exploratory ridge regression analysis, we asked: Is discrimination ability for stubby animate-looking objects associated with the ability to a) discriminate object-matched faces, b) discriminate self-matched faces, and c) recognize faces? We made aggregate dependent variables and chose ridge regression over linear regression to minimize possible effects of multicollinearity, as dependent variables in our preregistered analysis pipeline were highly correlated (see supplementary table s1; for corresponding analysis with linear regression, see supplementary table s7; see also additional analyses, including sensitivity analysis and models where the contribution of each quadrant of object space is estimated separately in supplementary tables s2-s7). Ridge regression is a method that can be used to fit a regression model when there is multicollinearity of the indicator variables, which can result in more accurate estimations of regression coefficients.

We performed three ridge regressions with demographics, control foraging, discrimination ability for stubby animate-looking objects, and discrimination ability for other objects as predictors. In all cases, discrimination ability for stubby animate-looking objects was an independent predictor of an individual’s face processing ability. Effect sizes were in all cases higher for stubby animate-looking compared to other objects (compare b-values in table 3) and including the former improved the ridge regression models by 1-2 percentage points. This further supports that individual differences in face processing are in part individual differences in the processing of stubby animate-looking objects.

**Table 3.**
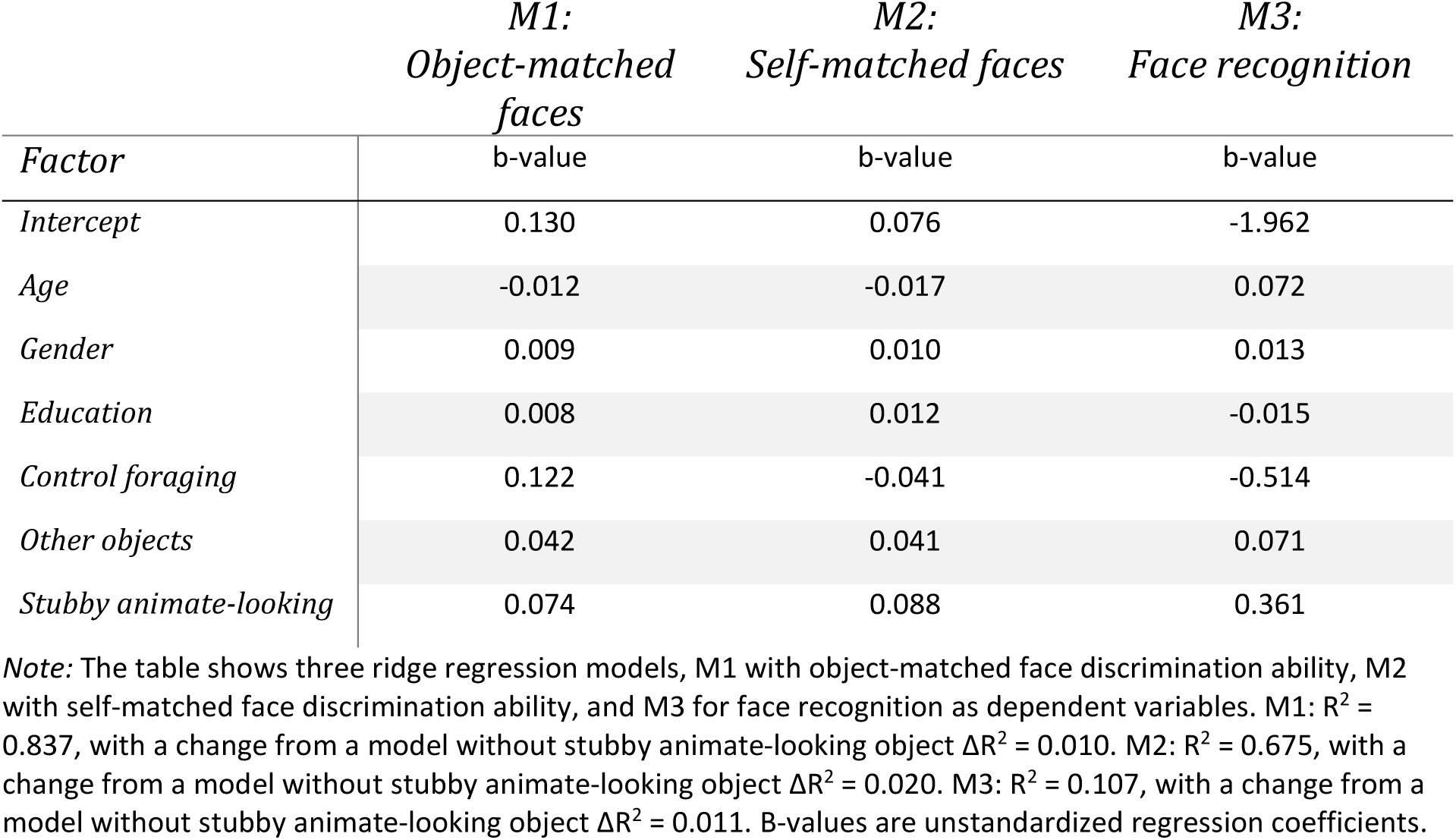
Predicting object-matched face discrimination, self-matched face discrimination and face recognition using ridge regression.

|  | *M1:*<br>*Object-matched*<br>*faces* | *M2:*<br>*Self-matched faces* | *M3:*<br>*Face recognition* |
| --- | --- | --- | --- |
| *Factor* | b-value | b-value | b-value |
| *Intercept* | 0.130 | 0.076 | -1.962 |
| *Age* | -0.012 | -0.017 | 0.072 |
| *Gender* | 0.009 | 0.010 | 0.013 |
| *Education* | 0.008 | 0.012 | -0.015 |
| *Control foraging* | 0.122 | -0.041 | -0.514 |
| *Other objects* | 0.042 | 0.041 | 0.071 |
| *Stubby animate-looking* | 0.074 | 0.088 | 0.361 |

*Note:* The table shows three ridge regression models, M1 with object-matched face discrimination ability, M2 with self-matched face discrimination ability, and M3 for face recognition as dependent variables. M1: $R^2 = 0.837$ , with a change from a model without stubby animate-looking object $\Delta R^2 = 0.010$ . M2: $R^2 = 0.675$ , with a change from a model without stubby animate-looking object $\Delta R^2 = 0.020$ . M3: $R^2 = 0.107$ , with a change from a model without stubby animate-looking object $\Delta R^2 = 0.011$ . B-values are unstandardized regression coefficients.

## Discussion

What are the diagnostic dimensions on which objects differ visually, and do these dimensions contribute to the organizational principles of visual object perception as evidenced by individual differences in behavior? Using data-driven methods, we utilized a deep convolutional neural network (Muttenthaler & Hebart, 2021; Simonyan & Zisserman, 2014) pretrained on ecologically relevant image categories (Mehrer et al., 2017; Mehrer et al., 2021) to construct a visual object space extracted from deep layer activations for a reference set of objects. The first two dimensions of object space capture attributes that distinguish one object from another and may map onto the anatomical organization of the ventral visual stream of non-human primates (Bao et al., 2020; but see Yao et al., 2023). Its existence in humans is unclear (Coggan & Tong, 2023; Yao et al., 2023; Yargholi & Op de Beeck, 2023).

We focused on the face quadrant of object space, occupied by stubby animate-looking objects such as faces. Faces evoke activity in specific ventral visual regions (Bao et al., 2020; Grill-Spector & Weiner, 2014; Kanwisher, 2000; Kanwisher et al., 1997), brain damage can lead to severe face recognition deficits (Davies-Thompson et al., 2014), and abilities in face processing vary greatly in the general population (Geskin & Behrmann, 2020; Ramon et al., 2019; Russell et al., 2009; Wilmer, 2017). As individual differences were crucial to our experimental design, we chose faces as our reference category and selected our sample to maximize such differences.

A significant proportion of face discrimination ability in our foraging task could be explained by demographics and performance on control trials, where older and less educated people in general were poorer at face discrimination than those younger and more educated, and those who did well on foraging for targets defined by simple features (black targets among white distractors or vice versa) tended to also be good at foraging for faces. These results may at least in part be due to various non-specific factors that correlate with one or more of these variables, such as motor control, attention, motivation, foraging organization (Ólafsdóttir et al., 2021), intelligence (Richler et al., 2017; 2019), and familiarity with computer-based tasks. Face discrimination ability was however additionally surprisingly well predicted by visual discrimination ability for miscellaneous other objects (compare to Richler et al., 2019), in accordance with individual differences in a general object perception factor *o* (Hendel et al., 2019; Richler et al., 2019).

When task was kept constant (foraging) and visual similarity of faces was kept comparable to those for non-face objects (object-matched face trials), demographics, performance on control trials, and performance on non-face object trials explained over 4/5^th^ of all variability in discrimination performance for faces – which is close to, but does not reach, the noise ceiling for face discrimination ability – and object performance explained a significant and large part of face discrimination ability not already accounted for by demographics and control performance.. It is possible that the discrepancy between our very strong association between object and face processing and some previous results on highly selective face processing abilities is task-dependent, as visual foraging is a reliable yet untraditional way to measure object discrimination ability. This discrepancy could also be experience-dependent, as recognition abilities for faces are more independent from general object recognition in people from larger hometowns (Sunday et al., 2019). The current study was done in Iceland, one of the smallest nations in the world, so the “face diet” in Iceland could be limited in comparison to more densely populated areas. Face processing in our sample could therefore to a larger extent tap into a general visual object perception ability *o*, or even an amodal domain-general object perception *o*-factor that may capture differences in the precision of working memory encoding across modalities (Chow et al., 2023). In general, our results provide strong support for the role of an *o*-factor in individual differences in visual object processing.

If the human visual system is however additionally organized around an object space, the processing of faces should partially depend on neural structures that also process other stubby animate-looking objects. Supporting this, we found association-dissociation patterns in individual differences in visual cognition where face processing was specifically associated with the ability to tell apart other stubby animate-looking objects. This contradicts previous work that found that visual similarity did not matter for individual differences in object recognition (Richler et al., 2019). The discrepancy may be due to task differences, greater variability in visual appearance of objects of the current study, or its unusually heterogeneous sample. Perhaps most likely, as the effect is small, it may only be found in very large-N studies and would likely be missed in any study that is not specifically designed to highlight it.

So far, three neuroimaging studies on a possible human object space have provided somewhat conflicting findings (Coggan & Tong, 2023; Yao et al., 2023; Yargholi & Op de Beeck, 2023). Using stimuli originally used to map object space in macaques (Bao et al., 2020), Coggan and Tong (2023) found extensive activations in the human ventral stream related to aspect ratio (stubby-spiky) and animacy. Yargholi and Op de Beeck (2023) on the other hand found little evidence for the human ventral stream being organized according to aspect ratio while anatomical organization was partially explained by animacy. While the study by Coggan and Tong (2023) may fit with the current results, activations in the human brain were not arranged in a clear sequence of complete maps of aspect-ratio/animacy object space as reported in non-human primates. Yao and colleagues (2023) also found that the topographic organization of the monkey brain is more regular than the human brain which they suggest may be due to distortion of the originally ordered topographic organization in the monkey brain during evolution. At a first glance, the study by Yargholi and Op de Beeck (2023) then appears to contradict the current study.

We believe that one likely explanation for the apparently diverging results has to do with effect size. While the effect related to the dimensions of object space that we see here is theoretically meaningful, it is small. To detect it, large samples are needed. Imaging studies usually cannot achieve this kind of power.

Another possible reason has to do with differences in the operational definitions of variables. We defined our object space by data-driven techniques using deep neural networks that capture image statistics of objects. Yargholi and Op de Beeck (2023) defined aspect ratio as a function of perimeter and area. As methodology was different, our results may not contradict those of Yargholi and Op de Beeck (2023) after all. Their stimuli were not linearly separable by aspect ratio in a 2D object space derived from the latent space of an artificial neural network, so is still an open question whether the human ventral stream is anatomically organized around the dimensions of object space as defined in such a way. These dimensions may or may not map directly onto *any* human labels, but covary with several of them, including animate-inanimate, stubby-spiky, and big-small real-world size (Konkle & Caramazza, 2013).

Another possible reason for discrepancies between studies is familiarity. In Coggan and Tong (2023), Yao et al. (2023), and Yargholi and Op de Beeck (2023), stimuli were primarily or exclusively familiar objects, making it hard to separate appearance from meaning and prior experience. Of note, these neuroimaging studies did not separate *being* animate/inanimate from *looking* animate/inanimate, and in Coggan and Tong (2023) all stubby animate-looking objects were human faces; Yao et al (2023) also localized stubby-animate-preferring areas using faces. Familiarity with objects may play a greater role when mapping out visual object space in humans compared to monkeys. Even though Bao et al. (2020) and (Coggan and Tong (2023) largely used the same stimuli, many of the things that have meaning to a human – such as a trumpet, camera, or a duck – may as well be “novel objects” for monkeys as they have no actual prior experience with them. Visual and semantic properties may both be represented in anterior regions of the human visual system (Proklova et al., 2016). Indeed, our results were particularly clear for novel objects as opposed to familiar objects. As experience with objects can shape the visual system (Sigurdardottir & Gauthier, 2015), the functional mapping of the anatomical organization of human object space may be clearer if based on objects with which people have no prior experience.

We suggest that the visual properties here approximated by two primary dimensions of object space are automatically extracted by the human visual system during an initial bottom-up sweep. These properties may guide the selection of appropriate methods of further object processing, as people are sensitive to various conceptual and perceptual properties of objects (Hebart et al., 2020). For example, the processing of animate objects may be skewed toward global, configural, or holistic processing (Maurer et al., 2002; Robbins & Coltheart, 2012a, 2012b) while inanimate objects could be preferentially processed in a piecemeal or feature-based manner (Peterson & Rhodes, 2003), and the processing of animate objects could additionally need greater integration across time due to their dynamic nature (Wallis & Bülthoff, 2001; see also Miyashita, 1993). Spiky objects could lend themselves to axis-based shape processing (shape skeletons; Blum, 1967; Firestone & Scholl, 2014; Hung et al., 2012; Kimia, 2003; Kovács et al., 1998; Kovács & Julesz, 1994; Lin, 1996; Sigurdardottir et al., 2014) or structural representations (parts and their relations; Biederman, 1985; Marr & Nishihara, 1978; Peissig & Tarr, 2007) while image-based or view-based representations (Peissig & Tarr, 2007; Riesenhuber & Poggio, 2000; Tarr & Bülthoff, 1998; Tarr & Pinker, 1989) may be more appropriate for stubby objects. At the same time, we emphasize that visual object processing may primarily utilize shared resources and mechanisms, as indicated by the substantial apparently domain-general individual differences in visual object discrimination abilities.

### Constraints on Generality

As recommended by Simons et al. (2017), we end with a Constraints on Generality (COG) statement aimed to identify and justify target populations – including people, situations, and stimuli – to pinpoint aspects believed to be crucial to observing the effect and those thought to be irrelevant. As people’s performance is affected by many non-specific factors that may vary from person to person, an important part of this project is the large sample size which allowed us to statistically partial out several such variables. Including several types of objects as stimuli also helps minimize systematic effects of varying experience (e.g., some people may be bird experts, but they are unlikely to only have expertise in stubby birds as opposed to spiky birds, and are furthermore unlikely to also be hand experts, bowtie experts, and seashell experts; expertise is therefore not confounded with stubby animate appearance). Only including target-distractor pairs within the same category (for familiar objects) also minimizes the role of semantic factors that may otherwise lead to noise and minimize effect sizes, as the semantic relationship between targets and distractors systematically affect object foraging (Ólafsdóttir et al., 2024).

Our study design required us to choose an object space model *a priori*, as we picked and matched our stimuli based on their coordinates in this space. We consider this a strength of our design as it allows precise experimental control of factors that might otherwise confound our results. We would however like to explicitly acknowledge that this means that the current study is based on this specific single instantiation of an object space model, while an infinite set of possible object space models exists. It is therefore possible that other variants of object spaces contribute even more – or do not contribute at all – to patterns of individual differences in object processing. Further, new data by Yao et al. (2023) suggests that high-level visual regions may be topographically organized around more dimensions than the two originally found by Bao et al. (2020). Likewise, our chosen object space, based on certain reference images processed through a specific layer of a particular neural network trained on a defined image set, is never going to be more than an approximation of any true object space of the human brain – which could partially explain our small effect size; any “true” effect of object space may therefore be underestimated.

We however do not think that our results should be limited to the specific reference image set chosen here, nor to the exact neural architecture or training set, as our view is that the two extracted dimensions are not arbitrary but reflect real-world image statistics. Bao et al. (2020; their extended data figure 4j) showed that projecting stimuli used to map the non-human primate anatomical object space to several network architectures resulted in a similar two-dimensional object space, so similar dimensions may be extracted from various neural networks. We note that prior to the current work, our smaller pilot study also showed that foraging for faces most greatly correlated with foraging for novel stubby animate-looking objects, but there we utilized a different network architecture (AlexNet layer fc6; Krizhevsky et al., 2012) and another image reference set (extended version of the Bank of Standardized Stimuli (BOSS; Brodeur et al., 2010, 2014) to define object space. We on purpose changed this in the current study to try to make sure that our results generalized and were not tied to a particular reference set or neural network architecture. Both the reference set and training set should however be varied and representative of real-world objects more generally, and our pilot work shows that unusual or homogenous reference sets (e.g., line drawings; images of text) can result in a two-dimensional object space that does not reflect dimensions closely related to animacy and aspect ratio.

Possible future replication studies should also be mindful to try to separate animate *appearance* (e.g., bowties, seashells, novel objects) from actually *being* animate objects; this was a particularly tricky part of the current study as these are highly correlated in real objects and are for some quadrants of object space impossible to fully dissociate – but it is in a minority of cases possible for the face quadrant. We focused on faces in the current study primarily to utilize well-documented individual differences in face processing abilities. A crucial aspect of our design was therefore to include people in our sample that maximized those differences – in other words to sample people ranging from severely impaired to excellent face recognizers.

This paper is, however, not intended to be only about faces, but about object perception more generally. Future studies should test whether our results generalize to other individual differences in object perception. For example, images of English and Chinese words are known to drive the visual word form area (VWFA) and project onto the two inanimate-looking quadrants of object space (Bao et al., 2020). Dyslexic readers have problems with recognizing visual words and are known to have a hypoactive VWFA (Richlan et al., 2011). Their visual recognition problems may generalize to some but not all objects (Jozranjbar et al., 2021; Kristjansson & Sigurdardottir, 2023; Sigurdardottir, Arnardottir, et al., 2021; Sigurdardottir et al., 2018; Sigurdardottir, Ólafsdóttir, et al., 2021). It would be highly interesting to test whether such problems would be exacerbated for inanimate-looking objects that share visual qualities with words.

We do not think that our results should be limited to object foraging. They may generalize to other tasks such as more traditional forced-choice recognition tasks or single-target visual search, especially if they cannot easily be solved via low-level visual strategies. Pilot studies however showed that compared to those tasks, object foraging had superior reliability and was additionally deemed the most enjoyable by participants. As the current study heavily relies on stable individual differences, possible replications of the current work need to take such factors into account.

Replications and extensions of the current work also need to consider our effect sizes – samples need to be large to capture small effects. Even larger samples than in the current study may be needed to fully compare our object space interpretation to other possible models explaining the data, such as through structural equation modeling (SEM). That said, small effects can still be theoretically meaningful; one white raven among a million black ravens proves that not all ravens are black. Our results support the contribution of object space to human visual cognition.

## Acknowledgements

We sincerely thank Sunneva Líf Albertsdóttir, Hannes Hansen, Hildur Franziska Hávarðardóttir, Martin Hebart, Unnur Andrea Ásgeirsdóttir, Ísak Örn Ívarsson, Isabel Gauthier, Bahareh Jozranjbar, Tim C. Kietzmann, Lukas Muttenthaler, Sarah Rutherford, Conor Smithson, Randi Starrfelt, Guðrún Þorláksdóttir, and all the people who shared images used in this study. This research was supported by the Icelandic Research Fund (grants no. 228916 and 218092) and the University of Iceland Research Fund.

## Author Note

### CRediT statement

H.M.S.: Conceptualization, methodology, software, formal analysis, resources, data curation, writing – original draft, writing – review & editing, visualization, supervision, project administration, funding acquisition.

I.M.O.: Conceptualization, methodology, software, formal analysis, data curation, writing – review & editing, supervision, project administration, funding acquisition.

We have no known conflict of interest to disclose. This study has been presented at conferences. This study was preregistered (https://osf.io/q5ne8). Data and analysis code are publicly available (https://osf.io/2jn6c).

## Supplementary Information

supplementary figure s1 shows the distribution of foraging ability for all foraging conditions and for face recognition ability.

**Supplementary figure s1.**
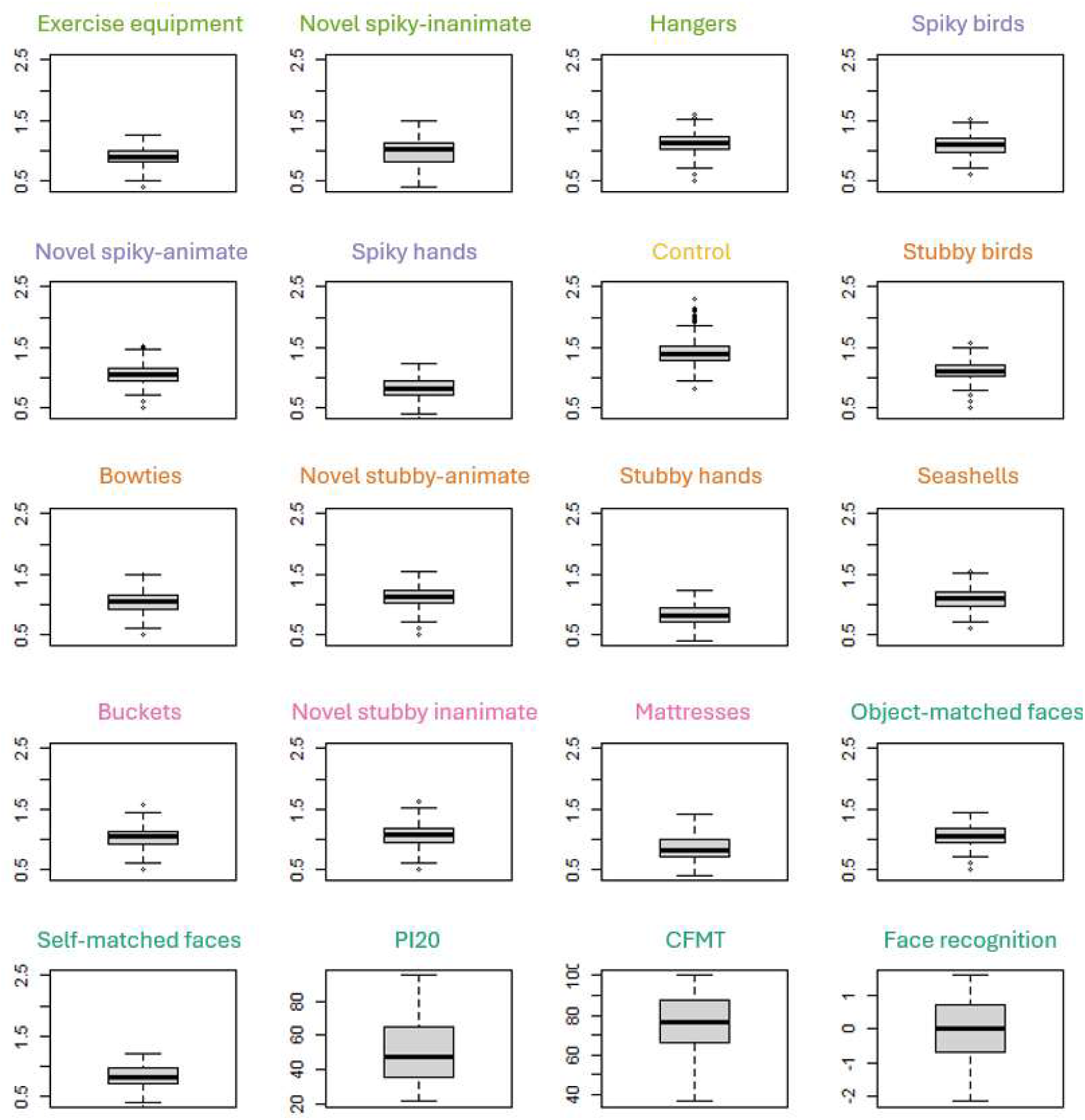
Boxplot showing the distribution of foraging and face recognition (N = 511). Note: Foraging performance is shown as median targets per second. 20-item prosopagnosia index (PI20) scores can range from 20 (no reported problems with faces) to 100 (most severe problems with faces). Cambridge Face Memory Test (CFMT) scores are in percent correct; chance level performance is 33%. Face recognition scores are in arbitrary units (scores from PI20 and CFMT were z-scored, the z-score of PI20 was negated, and the mean of these were taken).

supplementary figures s2 and s3 show foraging performance by age and education, respectively. Foraging performance decreased with age for all conditions and increased with educational level for all conditions except control trials.

**Supplementary figure s2.**
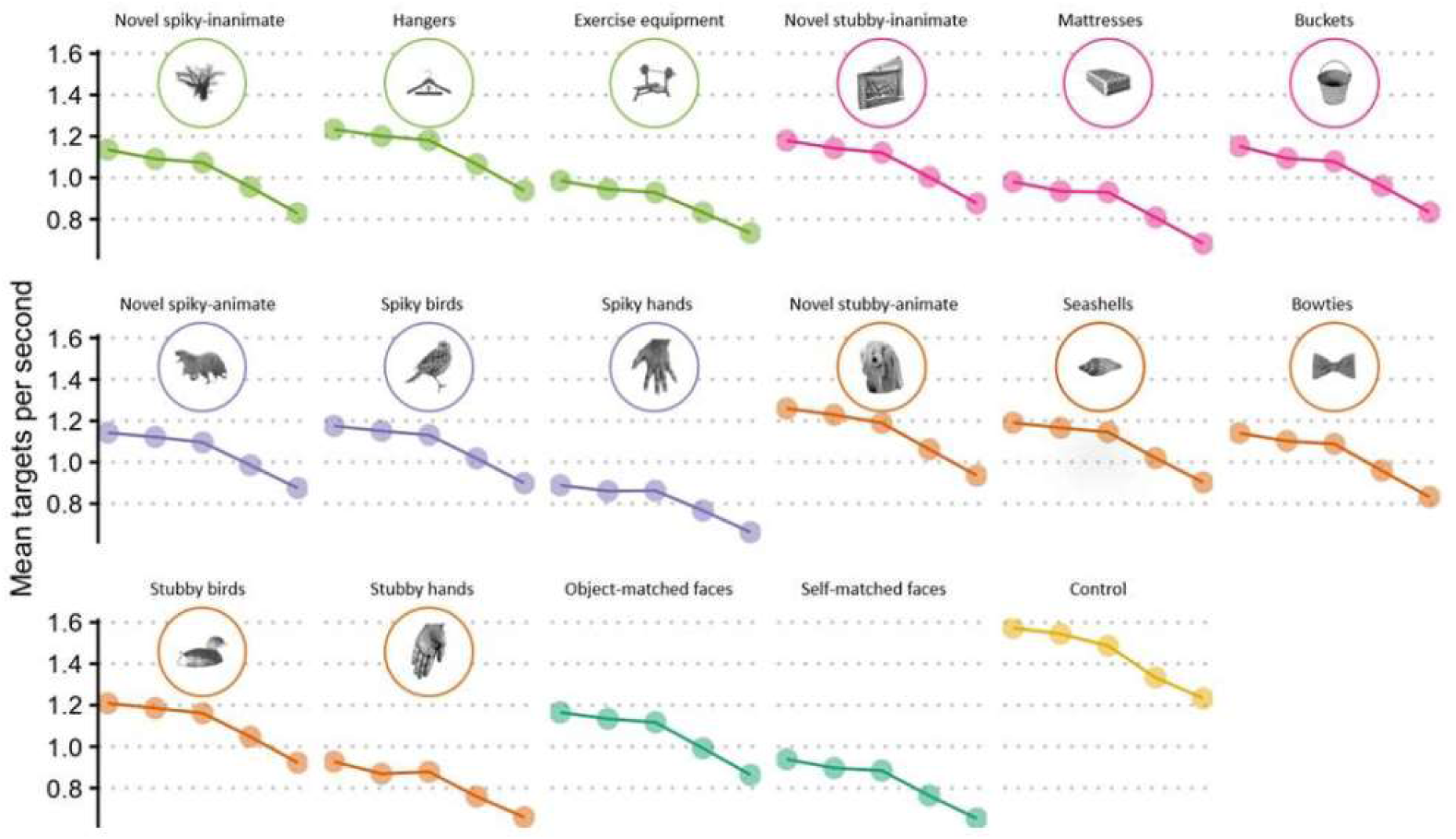
Foraging by age. Note: The five dots in each subplot represent the five age groups. Foraging performance declined with age for all trial types.

**Supplementary figure s3.**
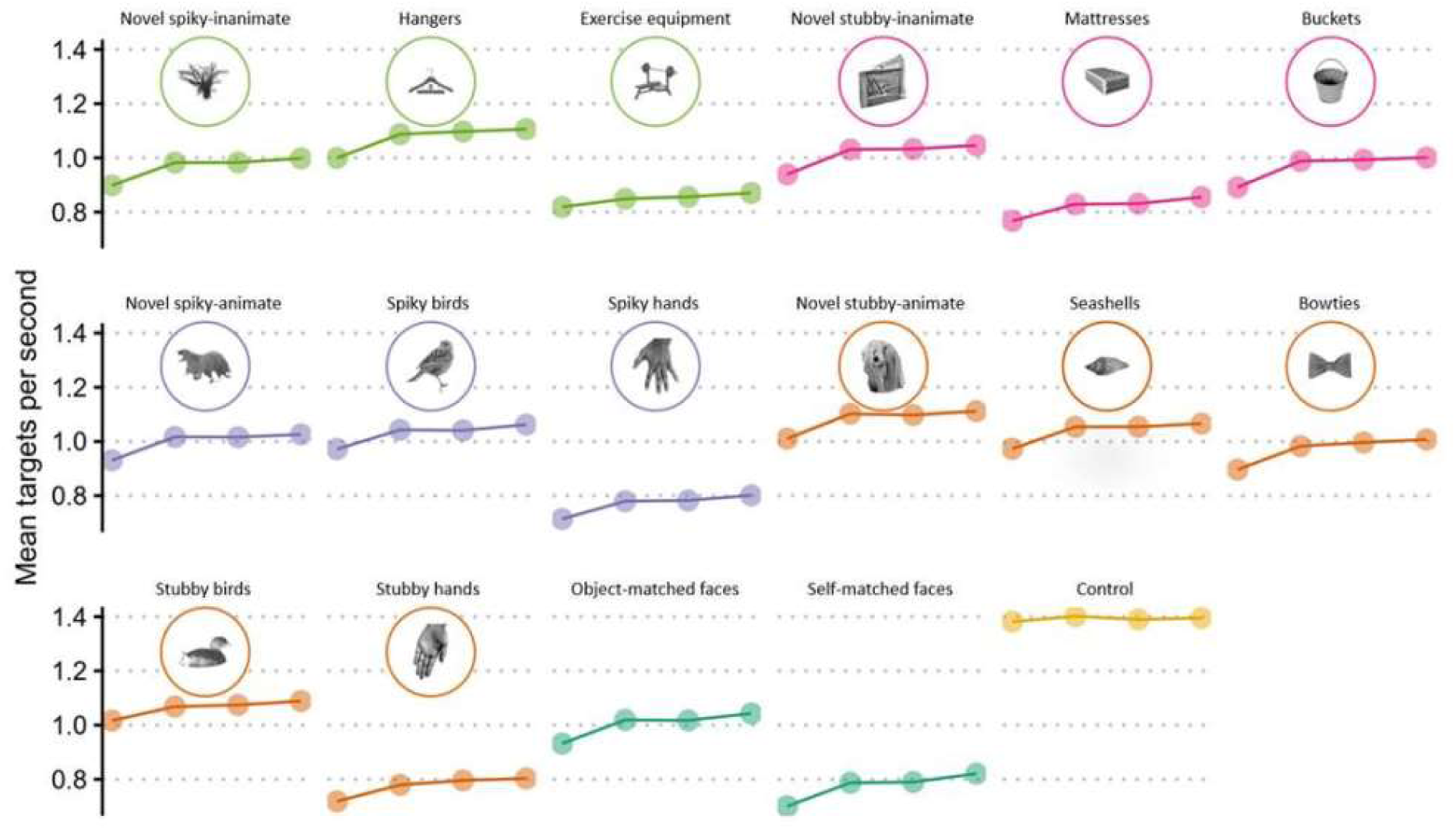
Foraging by education. Note: The four dots in each subplot represent the four educational levels. Foraging performance increased with educational level, with the notable exception of control foraging.

**Supplementary table s1.** Zero-order correlations between conditions.

|  | 1 | 2 | 3 | 4 | 5 | 6 | 7 | 8 | 9 | 10 | 11 | 12 | 13 | 14 | 15 | 16 | 17 | 18 |
| --- | --- | --- | --- | --- | --- | --- | --- | --- | --- | --- | --- | --- | --- | --- | --- | --- | --- | --- |
| 1 | -- | .83 | .82 | .83 | .82 | .78 | .67 | .83 | .81 | .82 | .77 | .82 | .83 | .83 | .79 | .79 | .72 | .23 |
| 2 |  | -- | .87 | .86 | .86 | .82 | .71 | .86 | .87 | .87 | .79 | .86 | .86 | .88 | .82 | .84 | .76 | .29 |
| 3 |  |  | -- | .88 | .90 | .77 | .80 | .90 | .89 | .91 | .78 | .91 | .89 | .90 | .82 | .87 | .73 | .25 |
| 4 |  |  |  | -- | .87 | .78 | .76 | .89 | .86 | .89 | .79 | .88 | .86 | .88 | .82 | .86 | .75 | .25 |
| 5 |  |  |  |  | -- | .78 | .77 | .88 | .86 | .90 | .78 | .88 | .86 | .89 | .81 | .87 | .76 | .29 |
| 6 |  |  |  |  |  | -- | .63 | .78 | .79 | .79 | .80 | .78 | .77 | .81 | .77 | .76 | .71 | .21 |
| 7 |  |  |  |  |  |  | -- | .81 | .74 | .82 | .58 | .78 | .73 | .76 | .65 | .77 | .60 | .16 |
| 8 |  |  |  |  |  |  |  | -- | .87 | .90 | .77 | .90 | .88 | .89 | .82 | .87 | .75 | .26 |
| 9 |  |  |  |  |  |  |  |  | -- | .89 | .81 | .89 | .88 | .88 | .84 | .86 | .78 | .29 |
| 10 |  |  |  |  |  |  |  |  |  | -- | .78 | .90 | .89 | .90 | .81 | .89 | .79 | .28 |
| 11 |  |  |  |  |  |  |  |  |  |  | -- | .81 | .79 | .81 | .79 | .77 | .75 | .27 |
| 12 |  |  |  |  |  |  |  |  |  |  |  | -- | .89 | .89 | .83 | .86 | .76 | .29 |
| 13 |  |  |  |  |  |  |  |  |  |  |  |  | -- | .89 | .83 | .86 | .77 | .30 |
| 14 |  |  |  |  |  |  |  |  |  |  |  |  |  | -- | .83 | .86 | .75 | .27 |
| 15 |  |  |  |  |  |  |  |  |  |  |  |  |  |  | -- | .80 | .75 | .28 |
| 16 |  |  |  |  |  |  |  |  |  |  |  |  |  |  |  | -- | .84 | .36 |
| 17 |  |  |  |  |  |  |  |  |  |  |  |  |  |  |  |  | -- | .46 |
| 18 |  |  |  |  |  |  |  |  |  |  |  |  |  |  |  |  |  | -- |

*Note:* 1. Spiky inanimate-looking exercise equipment; 2. Spiky inanimate-looking novel objects; 3. Spiky inanimate-looking hangers; 4. Spiky animate-looking birds; 5. Spiky animate-looking novel objects; 6. Spiky animate-looking hands; 7. Control foraging; 8. Stubby animate-looking birds; 9. Stubby animate-looking bowties; 10. Stubby animate-looking novel objects; 11. Stubby animate-looking hands; 12. Stubby animate-looking seashells; 13. Stubby inanimate-looking buckets; 14. Stubby inanimate-looking novel objects; 15. Stubby inanimate-looking mattresses; 16. Stubby animate-looking object-matched faces; 17. Stubby animate-looking self-matched faces; 18. Face recognition (aggregate of PI20 and CFMT).

**Supplementary table s2.** Association between face discrimination (object-matched faces) and discrimination of stubby animate-looking objects using ridge regression.

| *Factor* | *Hypothesis 1: Novel* | *Hypothesis 2: Familiar* | *Models* |
| --- | --- | --- | --- |
|  | b-value | b-value |  |
| *Intercept* | 0.142 | 0.105 | M1 |
| *Age* | -0.012 | -0.009 |  |
| *Gender* | 0.006 | 0.010 |  |
| *Education* | 0.009 | 0.005 |  |
| *Control foraging* | 0.094 | 0.088 |  |
| *Novel spiky-inanimate* | 0.143 | - |  |
| *Hanger spiky-inanimate* | - | 0.106 |  |
| *Exer. eq. spiky-inanimate* | - | 0.047 |  |
| *Novel stubby-inanimate* | 0.154 | - |  |
| *Bucket stubby-inanimate* | - | 0.106 |  |
| *Mattr. stubby-inanimate* | - | 0.052 |  |
| *Novel spiky-animate* | 0.208 | - |  |
| *Bird spiky-animate* | - | 0.118 |  |
| *Hand spiky-animate* | - | 0.040 |  |
| *Novel stubby-animate* | 0.234 | - | M1 & M2 |
| *Bird stubby-animate* | - | 0.111 |  |
| *Hand stubby-animate* | - | 0.074 |  |
| *Bowtie stubby-animate* | - | 0.083 |  |
| *Seashell stubby-animate* | - | 0.080 |  |

*Note:* The table shows ridge regression models for preregistered hypotheses 1 and 2. Dependent variable is face discrimination ability (object-matched faces). Regressions are run separately for novel objects and familiar objects. In both cases, M1 includes all factors except for foraging for stubby animate-looking objects, and M2 includes all factors, including foraging for stubby animate-looking objects. For novel objects, M2: $R^2 = 0.819$ , with a change from the M1 model without stubby animate-looking objects $\Delta R^2 = 0.010$ . For familiar objects, M2: $R^2 = 0.836$ with a change from the M1 model without stubby animate-looking objects $\Delta R^2 = 0.005$ . B-values are unstandardized regression coefficients.

A large part of the variance in face discrimination ability seems to be explained by background variables and a general object perception factor, with stubby animate-looking stimuli improving the model by about 1%. The added explanatory value of the model is significant, but not large. To determine whether this improvement of the model was due to the stimuli being stubby animate-looking or solely to the fact that we were adding another predictor to the model, we decided to redo our hierarchical regressions, adding performance for stimuli from three different quadrants of object space to the model in step 3, and adding performance for stimuli from the left-out quadrant in step 4. The results can be seen in supplementary table s3, which displays the change in R^2^ between steps 3 and 4 in the original hierarchical regressions, where performance for stubby animate-looking stimuli was added to the models in step 4, and the three alternative regressions where performance for each of the other quadrants was added at step 4. For both novel and familiar objects, adding the stubby animate-looking performance to the regression in step 4 has the largest effect on the explained variance of face discrimination abilities. Interestingly, for familiar objects, performance for spiky inanimate-looking objects does not significantly improve the model. The spiky inanimate-looking objects are the objects that are most different from faces, being neither stubby nor animate-looking.

**Supplementary table s3.** R^2^ change between step 3 and step 4 in hierarchical regression models.

| Quadrant of object space | $\Delta R^2$ novel objects | $\Delta R^2$ familiar objects |
| --- | --- | --- |
| Stubby animate-looking | 0.0103*** | 0.0064*** |
| Spiky animate-looking | 0.0073*** | 0.0038** |
| Stubby inanimate-looking | 0.0027** | 0.0035** |
| Spiky inanimate-looking | 0.0032** | 0.0018 |

*Note: \*\* $p < 0.01$ , \*\*\* $p < 0.001$ . This table displays the $\Delta R^2$ between steps 3 and 4 in hierarchical regression*
*models where the dependent variable is face discrimination ability, and stimuli from different quadrants of object space are added to the model in step 4.*

supplementary table s4 shows three regression models where 1) object-matched faces, 2) self-matched faces, and 3) face recognition are predicted by demographics, control foraging trials, and object foraging trials from each quadrant of object space separately. Interestingly, foraging for spiky inanimate-looking objects, which are the most dissimilar to stubby animate-looking objects, does not significantly improve any of the three models. In all three models, the stubby inanimate foraging trials have the lowest *p*-value and in two out of three models (object-matched faces and self-matched faces) they have the highest b-value of all quadrants.

**Supplementary table s4.** Predicting object-matched face discrimination, self-matched face discrimination and face recognition.

| *Factor* | *M1: Object-matched faces* |  |  | *M2: Self-matched faces* |  |  | *M3: Face recognition* |  |  |
| --- | --- | --- | --- | --- | --- | --- | --- | --- | --- |
|  | *b*-value | Std. Err. | *p*-value | *b*-value | Std. Err. | *p*-value | *b*-value | Std. Err. | *p*-value |
| *Intercept* | 0.091 | 0.035 | 0.009 | 0.041 | 0.049 | 0.404 | -1.970 | 0.406 | 1.66x10<sup>-6</sup> |
| *Age* | -0.009 | 0.003 | 0.006 | -0.015 | 0.005 | 0.002 | 0.089 | 0.039 | 0.024 |
| *Gender* | 0.011 | 0.007 | 0.126 | 0.013 | 0.010 | 0.188 | 0.024 | 0.084 | 0.774 |
| *Education* | 0.007 | 0.003 | 0.056 | 0.011 | 0.005 | 0.030 | -0.023 | 0.041 | 0.573 |
| *Control foraging* | 0.088 | 0.023 | 1.8x10<sup>-3</sup> | -0.094 | 0.033 | 0.005 | -0.687 | 0.274 | 0.012 |
| *Spiky inanimate-looking* | 0.030 | 0.024 | 0.213 | -0.021 | 0.035 | 0.550 | 0.223 | 0.286 | 0.436 |
| *Spiky animate-looking* | 0.064 | 0.024 | 0.007 | 0.058 | 0.034 | 0.089 | 0.285 | 0.278 | 0.307 |
| *Stubby inanimate-looking* | 0.049 | 0.023 | 0.035 | 0.058 | 0.033 | 0.077 | 0.545 | 0.271 | 0.045 |
| *Stubby animate-looking* | 0.081 | 0.018 | 6.2x10<sup>-6</sup> | 0.125 | 0.025 | 8.57x10<sup>-7</sup> | 0.507 | 0.207 | 0.015 |

We performed an additional sensitivity analysis (supplementary tables s5 and s6) where we split our large dataset into odd- and even-numbered participants and redid all analyses from Table 3 in the main manuscript with two independent datasets. In all cases, stubby animate-looking objects were significant independent predictors of face processing ability, and effect sizes (b-values) were in all cases higher for stubby animate-looking compared to other objects.

**Supplementary table s5.** Predicting object-matched face discrimination, self-matched face discrimination and face recognition for odd-numbered participants.

| *Factor* | *M1: Object-matched faces* |  |  | *M2: Self-matched faces* |  |  | *M3: Face recognition* |  |  |
| --- | --- | --- | --- | --- | --- | --- | --- | --- | --- |
|  | b-value | Std.E. | p-value | b-value | Std.E. | p-value | b-value | Std.E. | p-value |
| *Intercept* | 0.106 | 0.049 | 0.033 | 0.067 | 0.073 | 0.364 | -2.515 | 0.613 | 5.58x10<sup>-5</sup> |
| *Age* | -0.007 | 0.005 | 0.129 | -0.017 | 0.007 | 0.019 | 0.089 | 0.059 | 0.137 |
| *Gender* | 0.003 | 0.010 | 0.759 | 0.019 | 0.015 | 0.216 | 0.108 | 0.125 | 0.388 |
| *Education* | <0.001 | 0.005 | 0.944 | 0.016 | 0.007 | 0.026 | 0.033 | 0.058 | 0.574 |
| *Control foraging* | 0.066 | 0.029 | 0.022 | -0.082 | 0.043 | 0.058 | -0.453 | 0.358 | 0.207 |
| *Other objects* | 0.047 | 0.013 | 3.78x10<sup>-4</sup> | 0.043 | 0.020 | 0.031 | -0.018 | 0.164 | 0.911 |
| *Stubby animate-looking* | 0.087 | 0.025 | 6.32x10<sup>-4</sup> | 0.097 | 0.038 | 0.011 | 0.561 | 0.313 | 0.074 |

*Note: All reported p-values are two-sided, including for stubby animate-looking objects; a one-sided test (fitting a directional hypothesis of positive b-values) for stubby animate-looking objects would additionally be significant for face recognition.*

**Supplementary table s6.** Predicting object-matched face discrimination, self-matched face discrimination and face recognition for even-numbered participants.

|  | *M1: Object-matched faces* |  |  | *M2: Self-matched faces* |  |  | *M3: Face recognition* |  |  |
| --- | --- | --- | --- | --- | --- | --- | --- | --- | --- |
| *Factor* | b-value | Std.E. | p-value | b-value | Std.E. | p-value | b-value | Std.E. | p-value |
| *Intercept* | 0.071 | 0.049 | 0.152 | 0.010 | 0.068 | 0.884 | -1.423 | 0.549 | 0.010 |
| *Age* | -0.011 | 0.005 | 0.012 | -0.012 | 0.006 | 0.063 | 0.079 | 0.052 | 0.130 |
| *Gender* | 0.020 | 0.010 | 0.052 | 0.014 | 0.014 | 0.316 | -0.052 | 0.112 | 0.642 |
| *Education* | 0.015 | 0.005 | 0.005 | 0.006 | 0.007 | 0.395 | -0.083 | 0.058 | 0.151 |
| *Control foraging* | 0.126 | 0.039 | 0.001 | -0.123 | 0.053 | 0.020 | -1.283 | 0.430 | 0.003 |
| *Other objects* | 0.051 | 0.014 | $2.62 \times 10^{-4}$ | 0.021 | 0.019 | 0.265 | -0.014 | 0.154 | 0.928 |
| *Stubby animate-looking* | 0.065 | 0.025 | 0.011 | 0.157 | 0.035 | $8.38 \times 10^{-6}$ | 0.658 | 0.281 | 0.020 |

**Supplementary table s7.** Predicting object-matched face discrimination, self-matched face discrimination and face recognition using linear regression.

| *Factor* | *M1: Object-matched faces* |  |  | *M2: Self-matched faces* |  |  | *M3: Face recognition* |  |  |
| --- | --- | --- | --- | --- | --- | --- | --- | --- | --- |
|  | b-value | Std. Error | p-value | b-value | Std. Error | p-value | b-value | Std. Error | p-value |
| *Intercept* | 0.089 | 0.035 | 0.010 | 0.038 | 0.049 | 0.447 | -1.974 | 0.407 | 1.64x10<sup>-6</sup> |
| *Age* | -0.009 | 0.003 | 0.007 | -0.015 | 0.005 | 0.002 | 0.080 | 0.039 | 0.042 |
| *Gender* | 0.012 | 0.007 | 0.095 | 0.016 | 0.010 | 0.123 | 0.025 | 0.083 | 0.765 |
| *Education* | 0.007 | 0.003 | 0.047 | 0.011 | 0.005 | 0.021 | -0.021 | 0.041 | 0.610 |
| *Control foraging* | 0.088 | 0.023 | 1.69x10<sup>-4</sup> | -0.098 | 0.033 | 0.003 | -0.763 | 0.272 | 0.005 |
| *Other objects* | 0.048 | 0.009 | 5.67x10<sup>-7</sup> | 0.031 | 0.013 | 0.023 | 0.005 | 0.110 | 0.961 |
| *Stubby animate-looking* | 0.081 | 0.018 | 4.91x10<sup>-6</sup> | 0.128 | 0.025 | 4.55x10<sup>-7</sup> | 0.544 | 0.207 | 0.009 |

*Note:* The table shows three regression models, M1 with object-matched face discrimination ability, M2 with self-matched face discrimination ability and M3 for face recognition as dependent variables. M1: Residual standard error 0.070 on 504 degrees of freedom, $R^2 = 0.840$ , adjusted $R^2 = 0.838$ , F-statistic = 439.6 on 6 and 504 degrees of freedom, $p < 2.2 \times 10^{-16}$ . M2: Residual standard error = 0.100 on 504 degrees of freedom, $R^2 = 0.681$ , adjusted $R^2 = 0.677$ , F-statistic = 179 on 6 and 504 degrees of freedom, $p < 2.2 \times 10^{-16}$ . M3: Residual standard error = 0.826 on 504 degrees of freedom, $R^2 = 0.110$ , adjusted $R^2 = 0.099$ , F-statistic = 10.33 on 6 and 504 degrees of freedom, $p = 8.34 \times 10^{-11}$ . B-values are unstandardized regression coefficients.

